# Joint representation of working memory and uncertainty in human cortex

**DOI:** 10.1101/2021.04.05.438511

**Authors:** Hsin-Hung Li, Thomas C. Sprague, Aspen H. Yoo, Wei Ji Ma, Clayton E. Curtis

## Abstract

Neural representations of visual working memory (VWM) are noisy, and thus, decisions based on VWM are inevitably subject to uncertainty. However, the mechanisms by which the brain simultaneously represents the content and uncertainty of memory remain largely unknown. Here, inspired by the theory of probabilistic population codes, we test the hypothesis that the human brain represents an item maintained in VWM as a probability distribution over stimulus feature space, thereby capturing both its content and uncertainty. We used a neural generative model to decode probability distributions over memorized locations from fMRI activation patterns. We found that the mean of the probability distribution decoded from retinotopic cortical areas predicted memory reports on a trial-by-trial basis. Moreover, in several of the same mid-dorsal stream areas the spread of the distribution predicted subjective trial-by-trial uncertainty judgments. These results provide evidence that VWM content and uncertainty are jointly represented by probabilistic neural codes.

## Introduction

Working memory extends the duration over which neural representations are available to guide purposeful behaviors, and supports a wide range of cognitive functions, such as learning and decision-making ^1–3^. Although working memory is a fundamental building block of cognition, the neural activity that supports working memory is noisy and resource limited (reviewed in ^4^). Thus, decisions based on working memory are inevitably subject to uncertainty ^4–6^. Access to the uncertainty in our working memory enables us to use the extent to which we ‘trust’ our memory to make better decisions. Indeed, people’s reported confidence in their working memory performance correlates with the magnitude of memory errors, reflecting their ability to track the quality of their memory ^5,7–9^. Moreover, people incorporate knowledge of working memory uncertainty to improve their decisions in change detection tasks ^6,10,11^ and post-memory wagers ^12,13^.

Even though uncertainty plays a key role in supporting working memory-guided behavior, we know little about how working memory uncertainty is represented in the brain. Previous studies have established that the contents of visual working memory (VWM; e.g., the specific remembered orientation, color, motion direction, or spatial location) can be decoded from activation patterns in visual, parietal, frontal cortex and subcortical regions ^14–33^. However, these previous studies decoded VWM representations assuming a single point estimate of the memorized stimulus averaged over many trials. As we motivate next, ignoring both the distribution of decoded estimates and their trial-by-trial variability limits our ability to test theories of how neural populations encode VWM content and uncertainty, especially when it comes to links to memory behavior.

Neural population activity is noisy ^34–36^. According to the theory of probabilistic population codes, the brain knows the generative model that describes neural population activity as a function of stimulus features (e.g., location or orientation), including the distribution of the noise. Using this knowledge would make it possible to assess the appropriate level of uncertainty associated with a stimulus feature ^37–41^, a process known as ‘inverting’ the generative model. Under this theory, a population of neurons contains a joint representation of a stimulus along with uncertainty about the stimulus, and potentially even an entire Bayesian posterior probability distribution over the stimulus. Probabilistic population coding thus provides a testable hypothesis for how neural populations jointly represent a stimulus estimate and the associated uncertainty. In support of this hypothesis, previous studies reported that the probability distributions decoded from neural activity measured in visual cortex predict aspects of visual behavior ^42–44^. Here, we ask whether higher cognitive processes, like the items maintained by VWM, are also encoded as probability distributions by neural populations. We hypothesized that similar computational principles explain how neural populations maintain VWM representations. Specifically, we predicted that an item maintained in VWM is represented as a posterior probability distribution over the feature space (e.g., location). In this scenario, access to the content of VWM (e.g., remembered location) would involve a read-out of the mean of the distribution, while memory uncertainty would be reflected in the width of the distribution (**Fig. 1B**). The critical direct test of this hypothesis hinges on whether the parameters of the probability distribution actually predict the quality and uncertainty of measured memory behavior.

**Figure 1.**
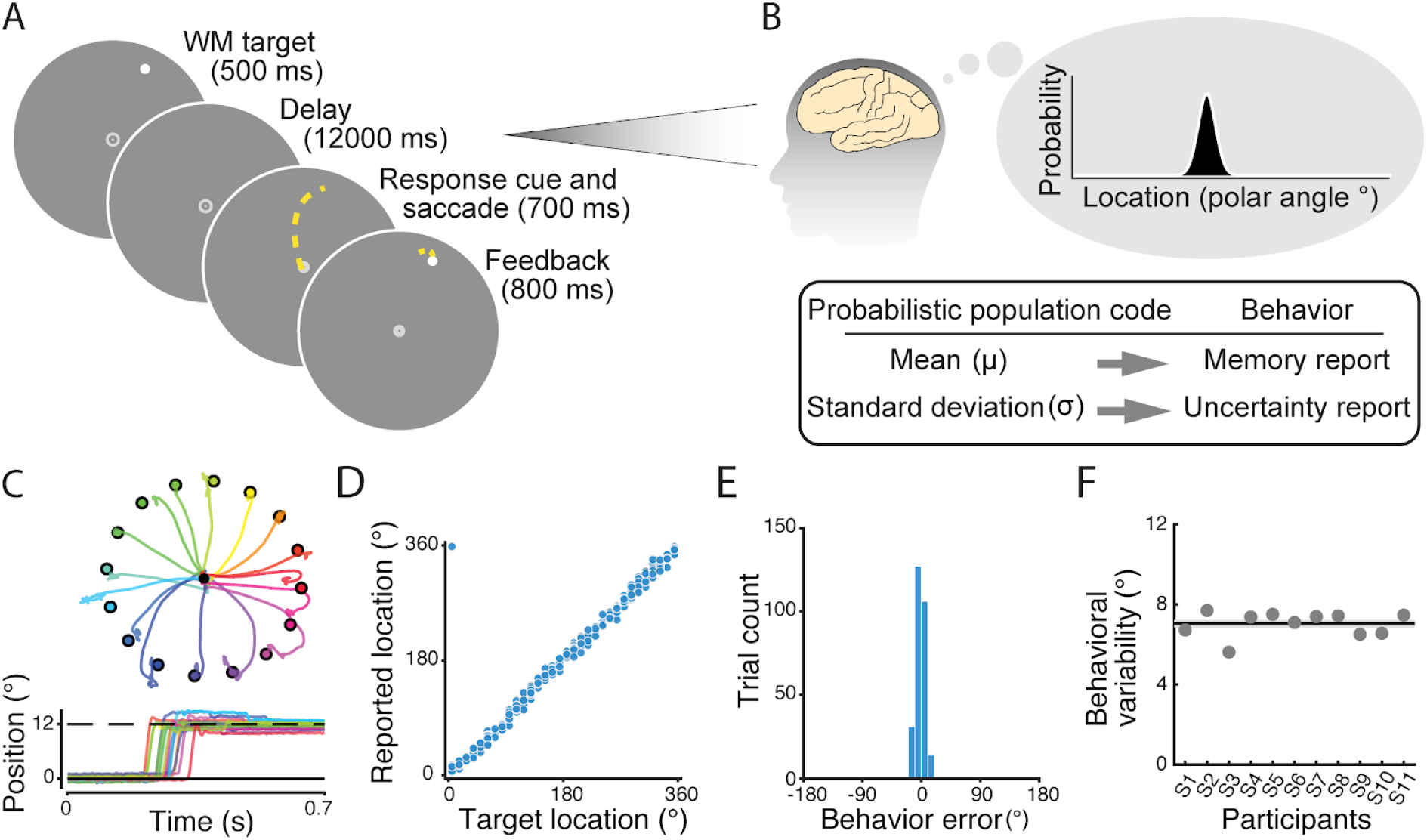
Procedures and working memory performance in Experiment 1. (A) Procedures. Participants maintained fixation while remembering the location of the target, presented at a pseudorandom position 12° from fixation. After which, participants generated a memory-guided saccade to the remembered location. Feedback, in the form of a white dot presented at the actual target location, permitted comparison with the landing spot following the memory-guided saccade. (B) We hypothesized that VWM is represented by a probabilistic population code. Per this hypothesis, populations of neurons represent the remembered target as a probability distribution over stimulus feature values (polar angle of the target in this case). This probability distribution allows a joint representation of the estimate of the memorized target (mean of the distribution) and the uncertainty of memory (standard deviation of the distribution). Two key predictions stem from this hypothesis: read-out of the mean of the maintained probability distribution guides memory reports, and the standard deviation of the probability distribution forms one’s memory uncertainty. (C) Example traces of memory-guided saccades for different locations across one scanning run (16 trials). The colored dots evenly spaced on an imaginary circle represent the target locations. (D) Memory reports from an example participant plotted against the target location. (E) Memory error distribution from the example participant in (D). (F) The variability of memory reports for individual participants (dots), quantified by the standard deviation of the memory error distribution. Dashed line shows mean across participants, and gray shaded interval shows ±SEM.

We measured fMRI BOLD activity in humans during two experiments to test each prediction from the above hypothesis. We then adapted and inverted a generative model for the BOLD activity ^42,45^. This yielded, on a trial-to-trial basis, a probability distribution over a memorized stimulus location from an activation pattern measured from retinotopic visual, parietal, and frontal cortex. While fMRI BOLD activity is subject to measurement noise, we still predicted that the decoded probability distribution would bear a resemblance to the one that the brain might use for its decision-making. In Experiment 1, we demonstrated that we can reliably decode the content of spatial VWM from BOLD activation. Moreover, trial-by-trial errors in the decoded positions predicted behavioral recall errors, revealing a close relationship between the decoded memory content and memory recall. In Experiment 2, we further established that the decoded uncertainty predicted explicit uncertainty reports when participants introspected the quality of their VWM in a wager task. Our results support the theory that the brain utilizes knowledge of the generative process of neural activity to represent memorized items probabilistically; in other words, that neural activity multiplexes the content of VWM and its uncertainty.

## Results

### Experiment 1

In Experiment 1, we used a Bayesian generative model to decode single-trial VWM representations from neural activation patterns, and assessed how the decoded memory content related to memory reports. We studied spatial VWM in a memory-guided saccade task. In each trial, we presented participants with a brief (500 ms) target dot, followed by a 12-second delay period (**Fig. 1A**). The polar angle of the target was chosen pseudo-randomly from 1 of 32 positions that spanned the full circle. Participants were asked to remember the location of the target while maintaining central fixation throughout the delay period. After the delay period, the empty fixation dot was replaced by a filled dot serving as the response cue. Upon the response cue onset, participants reported the remembered position by making a saccadic eye movement (e.g., ^46,47^; **Fig. 1A** and **Fig. 1B**). Behavioral memory reports were measured as the polar angle of the saccade endpoint.

We adapted a generative model ^42,45^ to decode a probability distribution over the stimulus location (polar angle) from the delay-period brain activity for each single trial. To focus on VWM maintenance activity, we used the averaged BOLD response for each voxel from 5.25 to 12.00 seconds after the delay period onset as the input to the model. The generative model described the multivariate voxel response given a stimulus location by a multivariate normal distribution. To estimate the mean of this distribution, the model approximated each voxel’s spatial tuning curve by a weighted sum of eight basis functions (channels) that evenly tiled visual (polar angle) space (**Fig. 2A**). For the covariance of the multivariate normal distribution, the model incorporated the empirical noise covariance estimated by the data and a theoretical noise covariance matrix that considered two sources of variability: the noise of each location channel and the noise specific to each voxel (see **Methods**). For each trial, we used the circular mean of the decoded probability distribution to represent the decoded remembered location.

**Figure 2.**
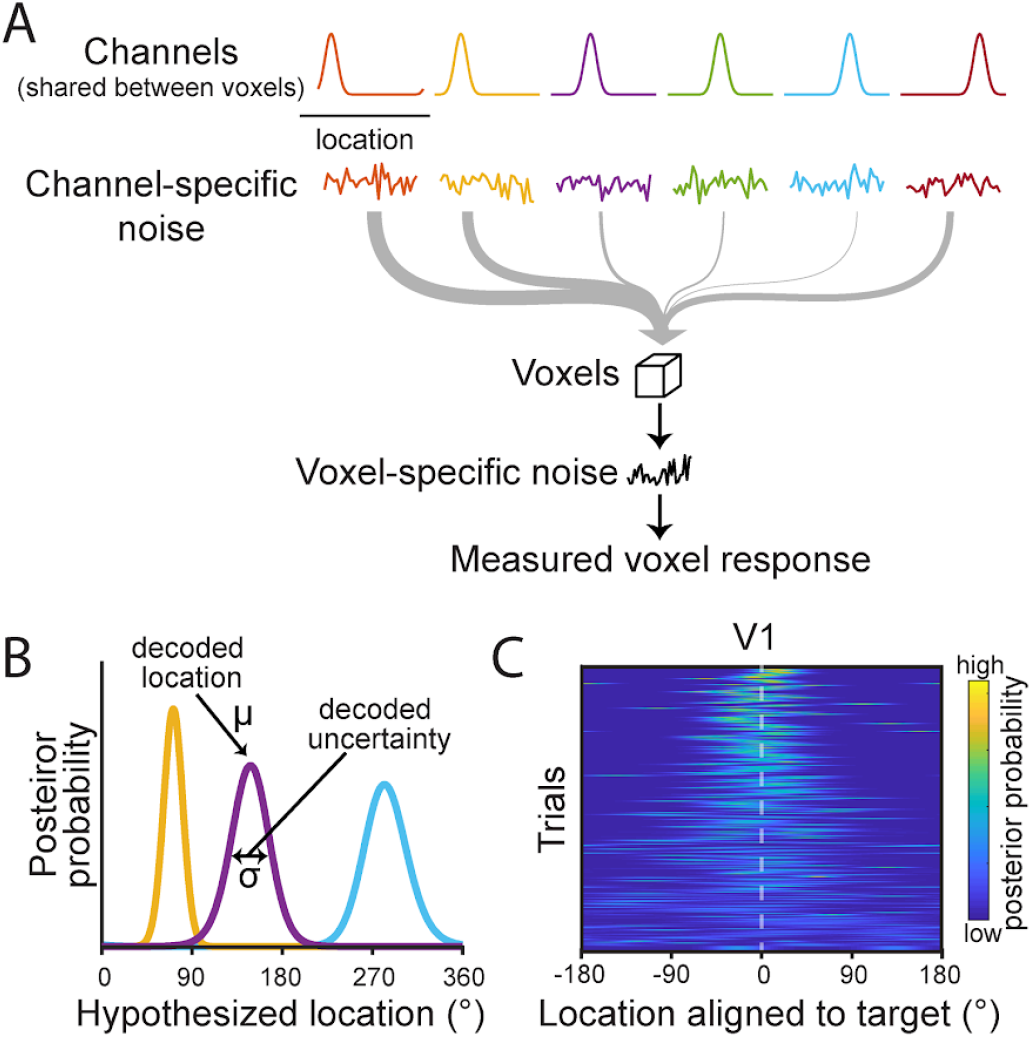
Generative model used to estimate and decode working memory representations. (A) Schematic of the generative model for BOLD response for spatial VWM (van Bergen & Jehee; 2021). The tuning function (mean response amplitude as a function of remembered target location) of each voxel is modeled as a weighted sum of eight basis functions evenly spanning the entire location space (0 to 360°; note: six shown here in cartoon depiction). Two sources of noise are considered: noise arising from each channel, which is shared across voxels, and noise arising from each voxel independently. (B) Posterior probability distributions decoded from the memory delay of three example trials. The decoded memorized location is derived from the circular mean of the posterior distribution. The decoded uncertainty in memory is derived from the circular standard deviation of the posterior distribution. (C) Posterior probability distributions decoded from an example participant’s primary visual cortex. Each row presents the posterior probability distribution decoded from the delay period of a single trial, where trials are sorted (from top to bottom) based on the decoded uncertainty of each trial (from lowest to highest uncertainty). The posterior distributions are circularly shifted to align to the target position of each trial (0°).

We first demonstrated that we can decode VWM content from delay period fMRI signals. We defined four retinotopic visual (V1, V2, V3 and V3AB), four parietal (IPS0, IPS1, IPS2 and IPS3) and two frontal areas (iPCS and sPCS) as regions of interest (ROIs) using population receptive field mapping techniques ^48,49^. Similar to previous studies using other decoding methods ^16,23,25,28,50^, we found that the remembered stimulus location could be decoded from the delay period BOLD responses in retinotopic visual, parietal, and frontal cortex. First, we plotted a distribution of the trial-wise decoding error (decoded location minus target location; **Fig. 3A**) for each ROI. These decoding error distributions reliably exhibited a single peak centered near 0° indicating the robustness of our decoder (**Supplementary Table 1**). We quantified the existence of decodable VWM information by comparing the standard deviation of the decoding error distributions to a null distribution obtained by a permutation procedure (see **Methods**). At the individual participant level, target locations were robustly decoded in most ROIs (**Supplementary Table 2**). At the group level, VWM contents were decoded in all ROIs (*p* < 0.001; unless otherwise noted, we report *p*-values corrected for multiple comparisons across ROIs via FDR with *q* = 0.05). The standard deviation of the decoding error distributions varied significantly across ROIs (permutation one-way repeated-measures ANOVA, *p* < 0.001, = 0.56; **Fig. 3B**), with smaller standard deviations in extrastriate cortex regions V3AB and V3, and IPS0 in intraparietal sulcus, indicating a more precise decoding performance in these regions. Two regions in the prefrontal cortex, iPCS and sPCS (the putative human homolog of macaque frontal eye field), had the largest standard deviations, indicating lower decoding performance in these areas. In another analysis, we obtained similar results when using the circular correlation between decoded location and target location to quantify the decoding performance (**Supplementary Fig. 1A** and **Supplementary Table. 3)**.

**Figure 3.**
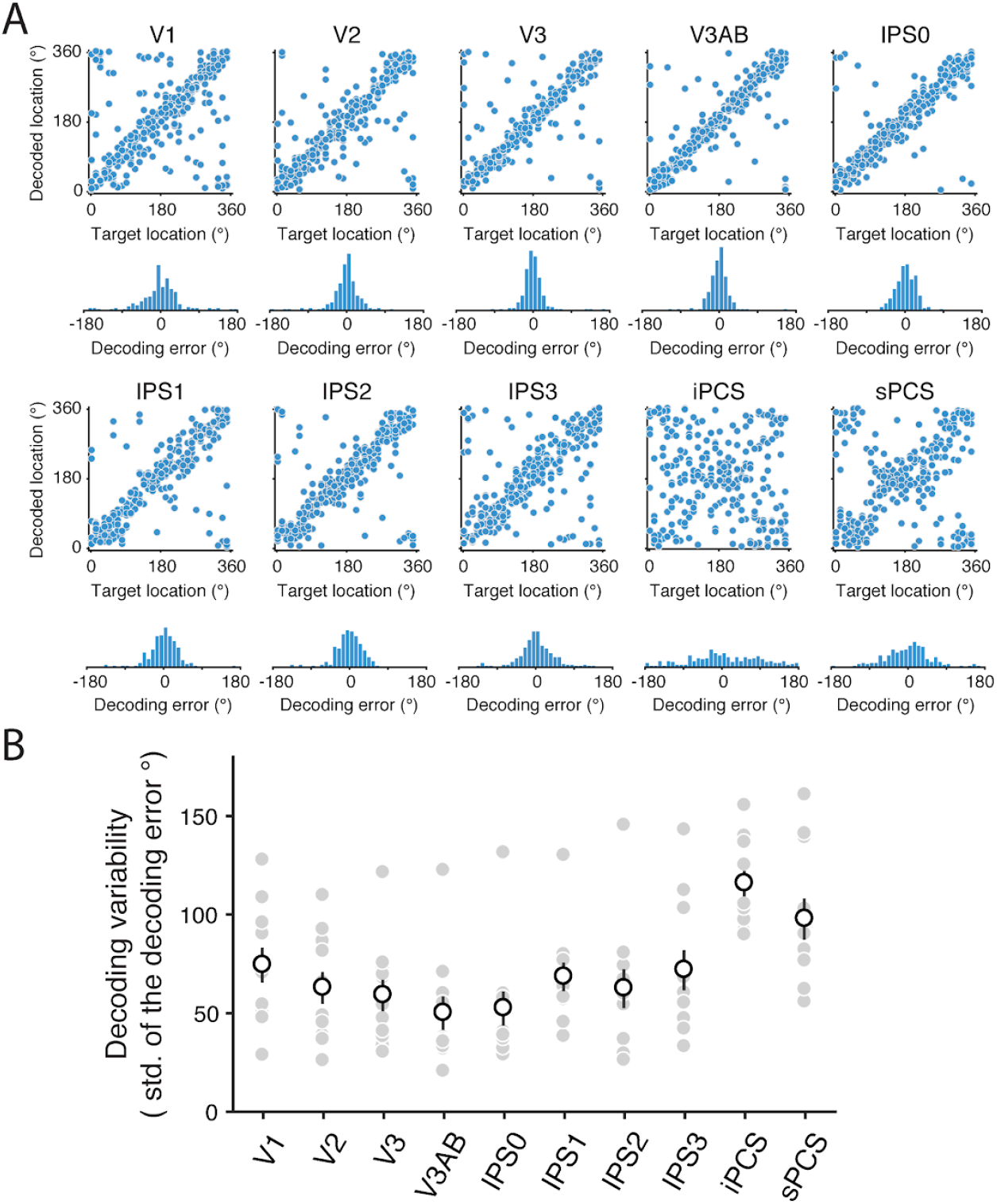
Working memory content can be precisely decoded. (A) Decoding performance of an example participant. For each ROI, the top figure represents the decoded location as a function of the memorized target location. The bottom figure is the distribution of signed decoding error (decoded location minus the memory target location). (B) Decoding performance quantified as decoding variability, the standard deviation of the decoding error distribution. The filled gray dots represent individual participants. The empty white dots represent the group average. The error bars represent ±SEM. Decoding performance varied significantly across ROIs (permutation one-way repeated-measures ANOVA, *F*(7, 70) = 12.6, *p* < 0.001,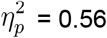).

As we are interested in working memory, we established that the signals we decoded cannot be attributed to sensory responses to the target stimuli. First, we are modeling the BOLD responses well into the delay period. Second, in a passive viewing experiment, a subset of participants (n = 3) performed a discrimination task at central fixation without the requirement to remember peripheral targets. Instead, we presented a highly salient, but irrelevant, flickering checkerboard at the same locations used in the WM task for the same duration as the VWM target stimulus (500 ms). Compared to the VWM experiment, the standard deviation of the decoding error distribution (averaged across subjects and ROIs) based on the same delay-period time points increased from 71° to 130° in the passive viewing experiment (**Supplementary Fig. 2A** and **2B)**. The near doubling of variability of the decoding error was only barely distinguishable from the null distribution in a subset of ROIs and participants (**Supplementary Table 2**). Moreover, the circular correlation between the decoded location and the target location was at zero under passive viewing for most participants and ROIs (**Supplementary Fig. 2C** and **Supplementary Table 3**). When we instead decoded stimulus location from earlier timepoints with strong evoked sensory signals (0.75 to 5.25 seconds from the delay deploy period onset), we were able to accurately decode stimulus location (**Supplementary Fig. 2D-2F**). Together, these results indicated that the neural representations of the target only persisted through the late delayed period when they were actively maintained in VWM.

So far, we have shown that we can decode the location of the memorized target. However, if the decoded VWM representation obtained from the BOLD signal drives behavioral performance, the decoded VWM representation should contain information relevant for behavioral memory reports beyond the physical location of the target. To investigate this issue, we leveraged our single-trial decoded locations, and we tested the prediction that (signed) memory error and (signed) decoding errors correlate at the trial-by-trial level. That is, we tested whether the direction of errors in memory and errors in decoding are the same (e.g., clockwise) with respect to the target. Accordingly, we computed the trial-wise circular correlation between memory error and decoding error for each participant and ROI. We found that the strength of this correlation varied across ROIs (permutation one-way repeated-measures ANOVA, *p* < 0.001, 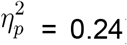). For individual ROIs, we found significant positive correlations in multiple regions including V2, V3, V3AB, IPS0, IPS2, IPS3 and sPCS (bootstrapping test, *p* < 0.05; **Fig. 4B**). Following previous studies using a similar Bayesian decoding approach ^42,45^, we quantified the correlations by binning the trials based on their decoding errors, computing the memory error of each bin and pooling the data across participants. We observed similar patterns as significant positive correlations were observed in multiple ROIs including V3, V3AB, IPS0, IPS2 (permutation test, *p* < 0.05; **Fig. 4C**). Overall, we found that memory behavior was linked to the neural representations we decoded, supporting our prediction that access to the content of VWM involves a read-out of the mean of the population-encoded probability distribution.

**Figure 4.**
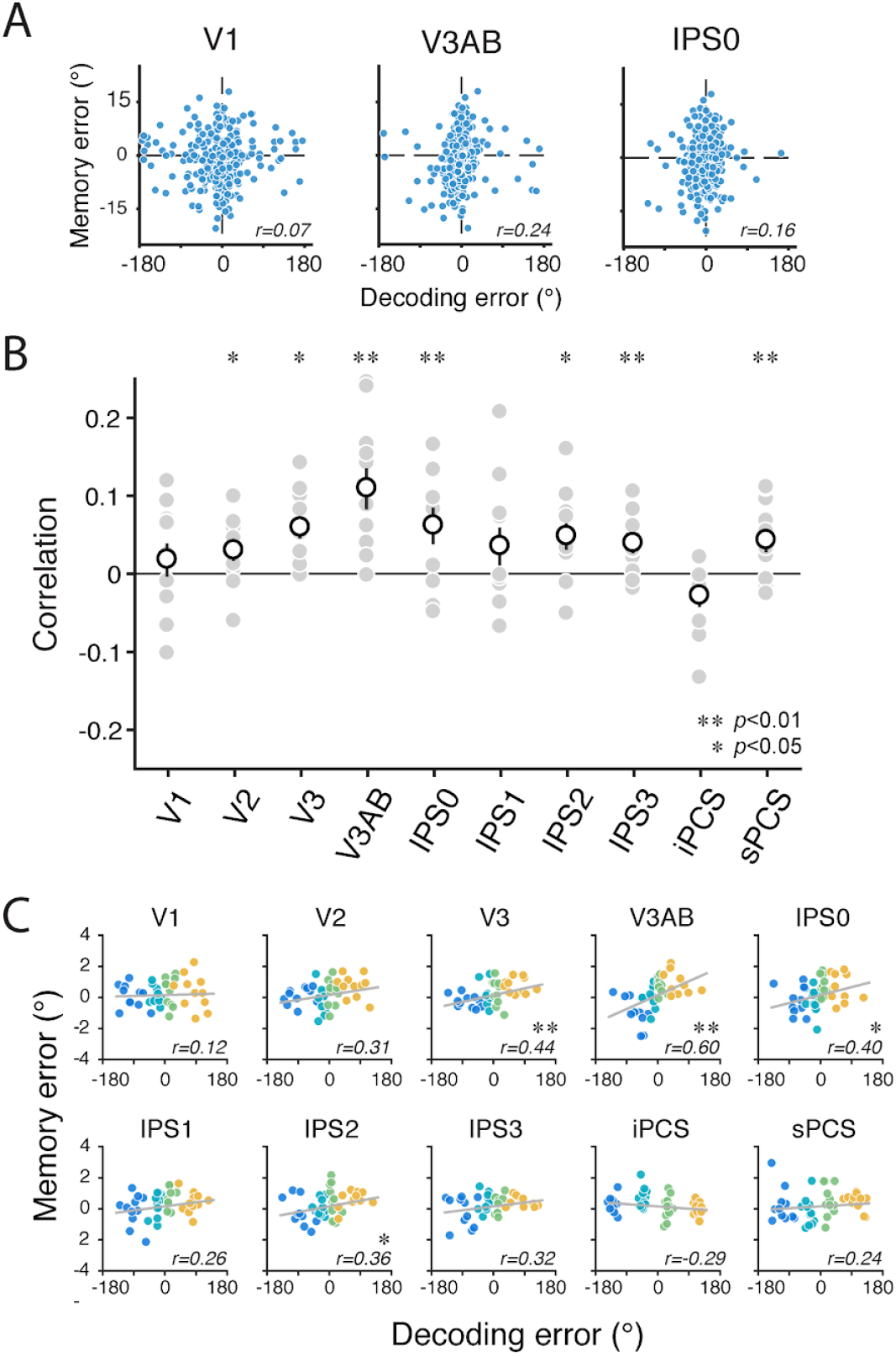
Errors in neural decoding of working memory predict behavioral memory errors in Experiment 1. (A) Memory errors plotted against neural decoding errors of an example participant for 3 ROIs. (B) Correlations are computed as circular correlations between the behavioral memory errors and neural decoding errors. The filled gray dots represent individual participants. The unfilled white dots represent the group average. The error bars represent ±SEM. **(C)** Memory errors plotted against decoding errors. The four colors indicate four bins (within each of 14 participants) sorted by decoding error. The gray line in each panel represents the best linear fit. The value at the lower right of each panel is the Pearson correlation coefficient.

### Experiment 2

In Experiment 1, participants reported the remembered location using a memory-guided saccade, and we quantified performance based on saccade landing position. We found that the population activity encoded not only the VWM target location, but additionally that errors in the decoder predicted errors in memory. Next, we tested the hypothesis that the population activity encodes a joint representation of both memory content and the uncertainty of their memory. Indeed, we can reflect on and directly report the confidence we have in our memory ^5,7,12,13^. Do these introspective reports reflect the uncertainty associated with the neural representation, quantified based on the posterior distribution decoded from neural activation patterns? To test this, in Experiment 2, we adapted our task so that participants were required to explicitly report the uncertainty of their memory with a wager.

The experimental procedures were similar to Experiment 1, with a few modifications. In addition to the filled dot at central fixation, the response cue contained a circular annulus with a radius matching the eccentricity of the target (**Fig. 5**). Participants reported their memory by making a saccade to the position on the annulus that matched the remembered target. Then, participants placed a wager by adjusting the length of an arc attached to the reported location ^12,13^. Participants were instructed to use the length of the arc to reflect the uncertainty of their memory. At the end of a trial, a feedback dot was presented at the true target location. Participants gained points if the target location was within the arc or otherwise gained zero points. The points they could gain decreased with the length of the arc, so observers were motivated to reflect their uncertainty using the arc length. In order to obtain the highest points, an optimal observer would increase the length of the arc with higher VWM uncertainty ^12,13^.

**Figure 5.**
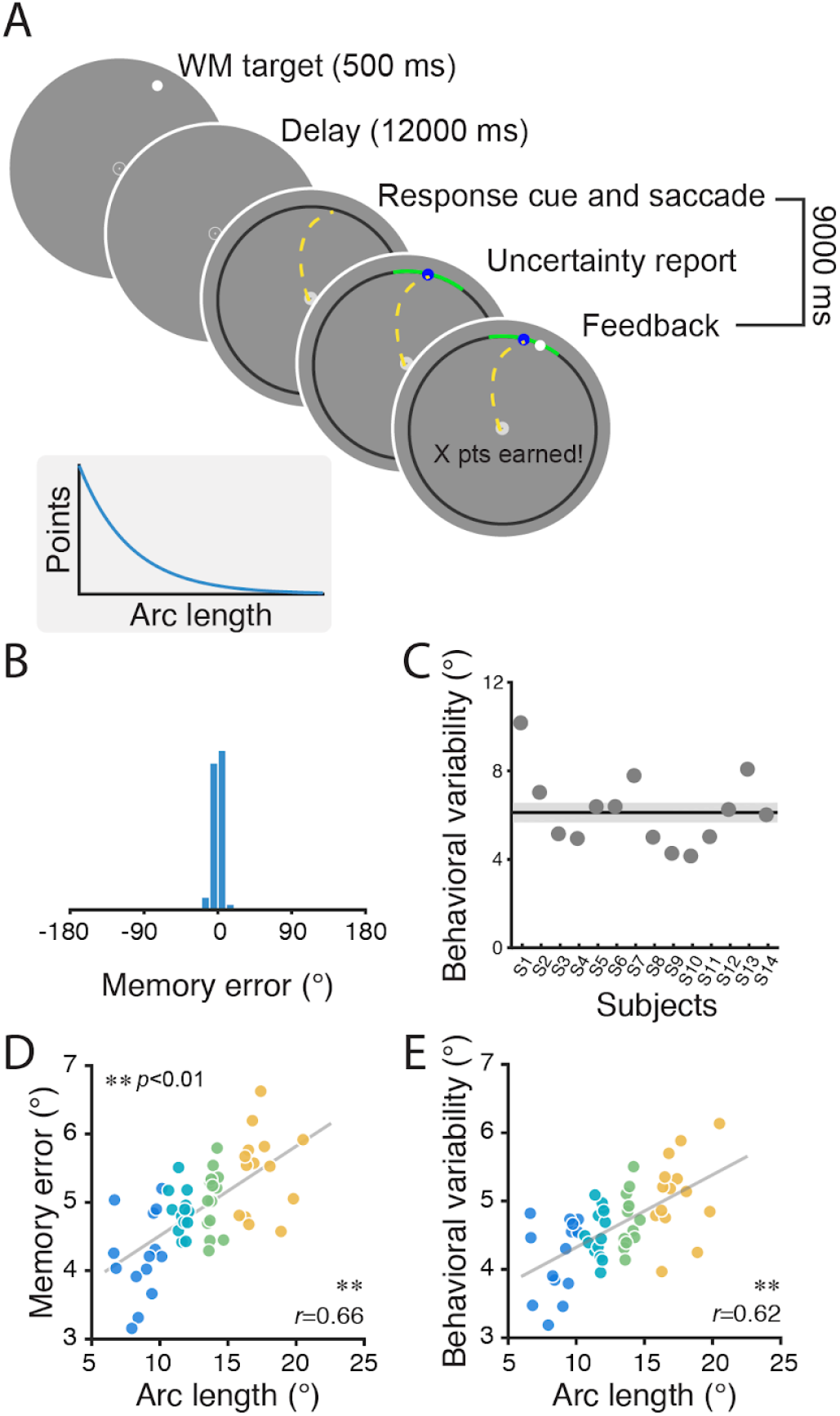
Procedures and working memory performance in Experiment 2. (A) The procedures were similar to those of Experiment 1 except for a few modifications. To report the remembered location, the participants generated memory-guided saccades onto a ring, and then reported their memory uncertainty by adjusting the length of an arc centered at the reported location. The trial ended with the onset of the feedback stimulus, a white dot presented at the target location. Participants earned points only if the target location was within the arc, and the points they earned decreased with the arc length. To earn a high score, participants should set shorter arcs when less uncertain. (B) The distribution of memory error from one example participant. (C) The variability of memory reports for individual participants, quantified by the standard deviation of the memory error distribution. (D) Memory error as a function of reported arc length, binned. Four colors represent four bins (within each of 14 participants) with increasing arc length. (E) Behavioral variability as a function of reported arc length. On trials where participants reported longer arc lengths, behavioral recall of remembered positions had larger errors (D; permutation test, p < 0.001) and was more variable (E; permutation test, p < 0.001). In (D) and (E) the gray lines represent the best linear fits.

Behaviorally, participants were able to monitor the quality of their VWM. Both the magnitude of memory errors (**Fig. 5D;** permutation test, *p* < 0.001) and the variability of memory reports (**Fig. 5E;** permutation test, *p* < 0.001) increased with the reported arc length. For some participants, the reported arc length varied as a function of target location, with shorter arc lengths and smaller errors at the cardinal angles (**Supplementary Fig. 3**). To evaluate whether the participants can track VWM uncertainty independent of the target location, we regressed-out the effect of target location (the polar angle between the target to the nearest cardinal angles) from the arc length. Still, the arc length correlated with the magnitude of memory error and the variability of memory reports (**Supplementary Fig. 4**), indicating that the participants’ ability to track the uncertainty across trials was not solely driven by the physical locations of the target. In sum, participants were not only aware of their memory uncertainty, but used these estimates to inform their wagers.

Next, we tested the hypothesis that these subjective estimates of memory uncertainty are jointly represented in the neural population activity that encodes the memory itself. To test this hypothesis, we correlated decoded uncertainty (standard deviation of the decoded posterior probability distribution) with behaviorally reported memory uncertainty (arc length, with the effect of target angle regressed out) at a single-trial level for each participant and each ROI. Decoded uncertainty correlated with the reported arc length significantly in V2, V3AB, IPS0 and IPS1 (bootstrapping test, *p* < 0.05; **Fig. 6B**). Additionally, we binned each participant’s trials based on decoded uncertainty and pooled the data across participants. Consistently, we found that participants reported larger arc length in trials with higher decoded uncertainty in V2, V3AB, IPS0 and IPS1 (permutation test, *p* < 0.05; **Fig. 6B**). These results support the notion that the uncertainty of VWM can be represented along with the memorized location by a probabilistic population code, and the uncertainty encoded in the neural population is used for explicit uncertainty reports.

**Figure 6.**
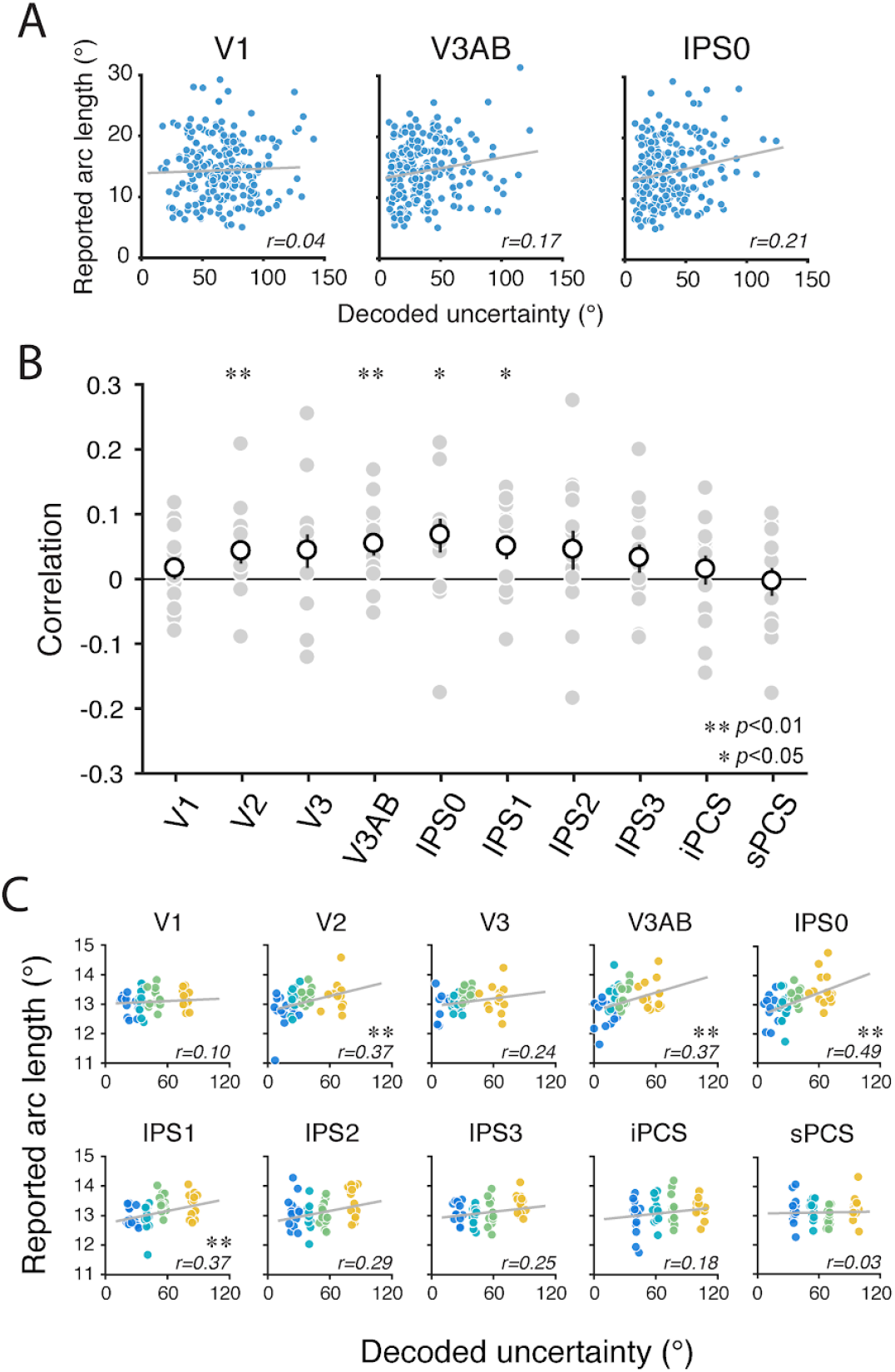
Decoded memory uncertainty predicts subjective memory uncertainty. (A) Reported arc length (subjective VWM uncertainty report) plotted against decoded uncertainty of an example participant for 3 ROIs. (B) Correlations between reported arc length and decoded uncertainty. The filled gray dots represent individual participants. The empty white dots represent the group average. The error bars represent ±SEM. (C) Reported arc length plotted against decoded uncertainty. The four colors indicate four bins (within each of 14 participants) with increasing decoded uncertainty. The gray line in each panel represents the best linear fit. The value at the lower right of each panel is the Pearson correlation coefficient.

In perceptual decision-making, people utilize their knowledge of their own reaction time when making uncertainty judgements ^51^. Thereby, saccade reaction time might implicitly track VWM uncertainty in both experiments. Behaviorally, reported arc length increased with saccade reaction time, indicating an impact of reaction time on uncertainty judgement (**Supplementary Figure 5A**). In terms of fMRI BOLD activity, saccade reaction time correlated with decoded uncertainty in V3AB and IPS0, when binning trials based on decoded uncertainty (**Supplementary Figure 5C** and **5E**).

Regarding the decoded memorized location, the results of Experiment 2 replicate those of Experiment 1. VWM contents were decodable in all the ROIs (permutation test, *p* < 0.001 for all ROIs). The precision of the neural decoding error distribution varied across ROIs (permutation one-way repeated-measures ANOVA, *p* < 0.001, 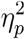; **Supplementary Fig. 6;** also see **Supplementary Table 4 - 6**), with the highest precision observed in V3AB and the lowest precision in iPCS and sPCS (**Supplementary Fig. 6**). To further evaluate the behavioral relevancy of the decodable information, we correlated (signed) memory error with (signed) decoding error. The main effect of ROI on this correlation was significant (permuted one-way repeated-measures ANOVA, *p* < 0.01, 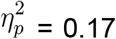). Memory error correlated with the neural error in all the ROIs, except iPCS (bootstrapping test, *p* < 0.05; **Fig. 7B**). We obtained similar results when binning each participant’s trials based on decoding error and pooled the data across participants (**Fig. 7C**).

**Figure 7.**
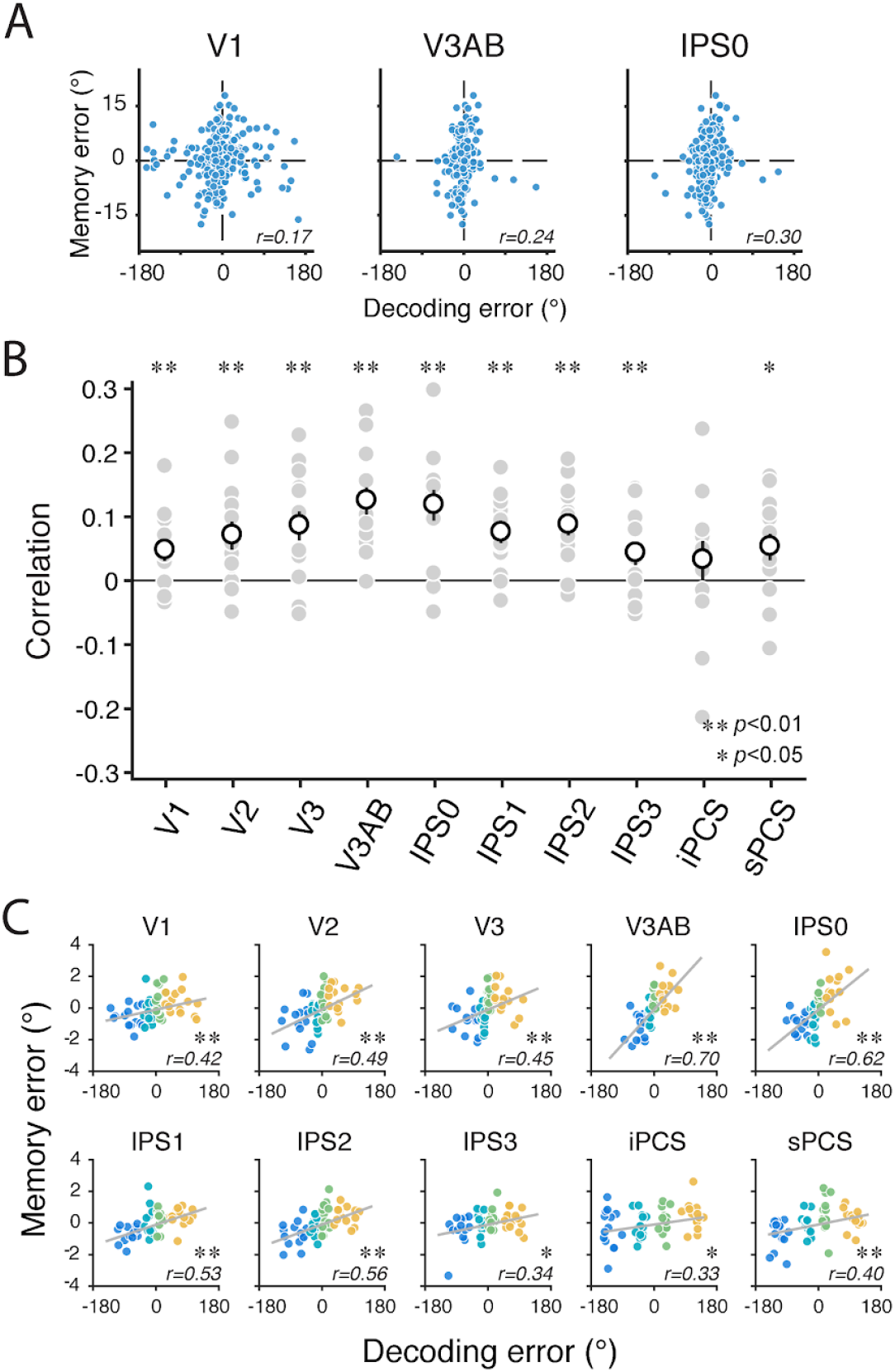
Errors in neural decoding of working memory predict behavioral memory errors in Experiment 2. (A) Behavioral memory error plotted against neural decoding error of an example participant. (B) Correlations are computed as circular correlations between the memory error and neural decoding error. The filled gray dots represent individual participants. The empty white dots represent the group average. The error bars represent ±1 SEM. (C) Memory error plotted against decoding error. The four colors indicate four bins (within each of 14 participants) sorted by the decoding error. The gray line in each panel represents the best linear fit. The value at the lower right of each panel is the Pearson correlation coefficient. Overall, these results replicate those reported in Experiment 1 (Figure 4).

While our model made direct predictions that the decoded location and decoded uncertainty are reflected in memory reports and uncertainty judgments, respectively, there could additionally exist a relationship between the decoded uncertainty and the variability of memory reports ^42^. Within the context of spatial VWM, decoded uncertainty did not correlate with the magnitude of memory error, or with the variability of memory reports (**Supplementary Fig. 7**). To investigate whether such relationships between decoded uncertainty and memory errors exist at a cross-subject level, for each participant, we averaged the decoded uncertainty across trials. Widespread across multiple ROIs in visual cortex and IPS, we found that participants with larger averaged decoded uncertainty performed worse in the behavioral memory reports when quantified as their averaged magnitude of behavioral memory error (**Fig. 8**) or as the standard deviation of their behavioral error distribution (**Supplementary Fig. 8**). These results demonstrate a linkage between the precision of VWM neural representation in these brain regions and the precision of behavioral memory reports.

**Figure 8.**
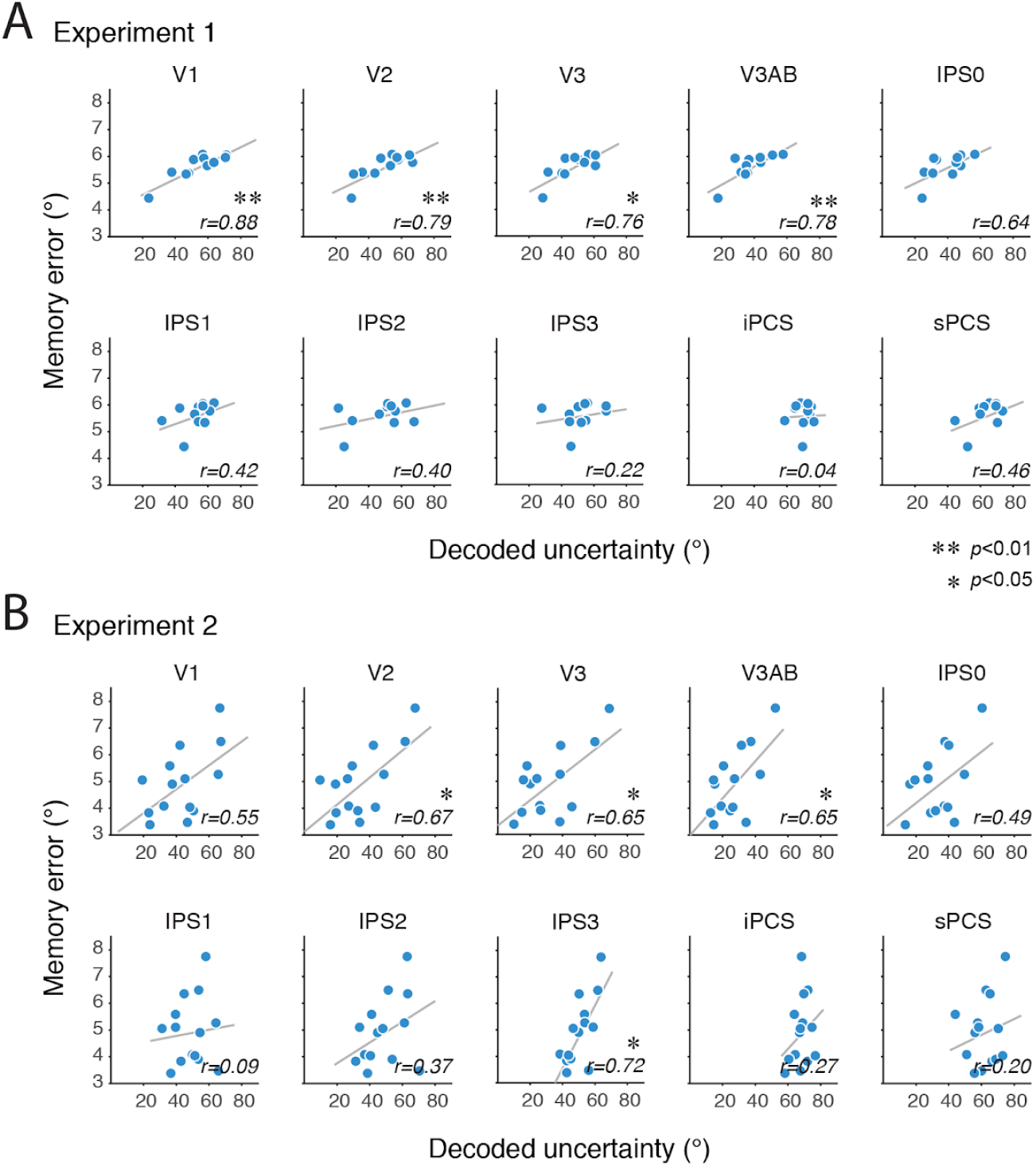
Participants with overall greater decoded uncertainty have less precise working memory. (A) Experiment 1. Each dot represents one participant. The decoded uncertainty (x-axis) and the absolute value of memory error (y-axis) was averaged across trials for each participant. The gray lines represent the best linear fit. (B) Experiment 2.

## Discussion

Although it is well established that the contents of working memory can be decoded from human brain activity, it remains unknown whether and how memory uncertainty is represented in the brain. Here, inspired by the theory of probabilistic population codes ^37,38,40,41^, we tested the hypothesis that the human brain encodes VWM as a probability distribution over the remembered feature space. In two independent experiments using a generative model of neural activity combined with a multivariate Bayesian decoder, we tested two central predictions that stem from this hypothesis. First, after validating that our procedures could decode the precise contents of VWM during a memory retention interval, we discovered that errors in our neural decoder predicted the direction and amplitude of memory errors made later in the trial. Second, we discovered that the uncertainty in our neural decoder predicted the memory uncertainty explicitly reported by our participants. Together, these results provide strong evidence that the content of our working memory is a read-out of a noisy probability distribution encoded in the population activity of neurons whose distribution width conveys information about memory uncertainty.

The theory of probabilistic population codes was originally proposed to explain how the brain can jointly represent the estimate and the uncertainty of sensory stimuli used during perception. Per this theory, the brain possesses knowledge about the generative process of neural activity ^37–41,52^. This knowledge and the variability in the response of cortical neurons naturally allow the neural population to represent the probability distribution over the perceptual stimulus space for any pattern of neural activity. Neurophysiological experiments measuring population-level neural responses have found evidence supporting the predictions of probabilistic population codes for visual perception in non-human primates ^44,53,54^. Here, we demonstrate that probabilistic population codes are not limited to visual perception, but are also used to represent information actively maintained in working memory in support of higher cognition. Our computational neuroimaging approach—utilizing the knowledge of a generative model and applying Bayesian decoding on the observed fMRI activation—mimics how the ‘decision maker’ in the brain performs inference based on its knowledge of its own generative model and the observed neural activity during the VWM delay period.

Under the rubric of probabilistic population codes, the fluctuations of both the content and the uncertainty of VWM arise from the noise in the neural population response that encodes the memorized stimulus. Thus, it is critical that we are able to decode VWM content and its uncertainty on a trial-by-trial basis in order to study their relationships with behavior. Previous neuroimaging studies have mostly reported VWM decoding accuracy (or fidelity) per condition, averaged across all trials in the experiment (e.g., ^14–26,28–30,55^). These indices represent decoding quality aggregated across many trials, and thus they are inadequate to explain or to estimate how VWM content and its uncertainty fluctuate across individual trials.

Unlike previous studies that used simpler linear encoding models ^56^ to decode the content of spatial VWM ^16,23,25,50^, we used a generative model that improves the precision of decoding by estimating sources of measurement and neural noise ^45^. In both experiments, we found remarkably precise representations of the memorized target locations in a widely distributed network of brain regions including visual, parietal, and frontal cortex. Encouraged by the robustness of the decoding, we asked whether these population responses in fact encoded the memory, including small spatial errors in memory, rather than the veridical target locations. On a trial-by-trial basis, we found that errors in our neural decoder predicted the direction and amplitude of memory errors (**Fig. 4**). These results indicate that the neural representations we decoded from the late delay period preceding the participants’ memory-guided responses, contain information that affects behaviours beyond that present in the physical stimulus. Specifically, it strongly suggests that one’s memory depends on the read-out of these population encoded representations. In neurophysiological studies, this type of correlation (between neural noise and behavioral choices) has sometimes been used to infer a causal link between neural responses and behaviors (reviewed in ^57^) We observed the strongest correlations in V3AB in dorsal extrastriate cortex, followed by neighboring regions V3 and IPS0. These results are generally consistent with neurophysiological studies of perceptual decisions reporting that choice-related activity is weak at best in early sensory cortex, and stronger in higher-tier sensory cortices ^57–63^.

Next, we leveraged the signal-trial decoding to investigate the neural basis of VWM uncertainty. As VWM uncertainty is defined as the width of one’s belief distribution over possible stimulus values (memorized location) or the subjective sense of the quality of one’s own memory, uncertainty can fluctuate across trials even when remembering the exact same stimulus. In line with previous behavioral studies ^5,7,9,12,13^, we found that uncertainty judgements tracked the quality of VWM on a trial-by-trial basis (**Fig. 5D** and **5E**). Importantly, this demonstrates that participants were aware of the quality of their memory and adjusted their uncertainty reports in step with their memory fidelity. We also found that the measured population encoded responses, when analyzed with our generative model, stored memory uncertainty. On a trial-by-trial basis, memory uncertainty decoded from the retention interval in V2, V3AB and IPS0 predicted the uncertainty explicitly reported and utilized by the participants in the wagers made later in the trial (**Fig. 6**). Recall we also observed strong correlations between decoding error and memory error in V3AB and IPS0. Theoretically, an estimate of an item and the uncertainty of the estimate can be jointly encoded as a single probability distribution by the same population of neurons. Our findings suggest that such an efficient mechanism exists in V3AB and IPS0 to support VWM.

Our theory-guided approach of decoding uncertainty from the width of a modeled probability distribution is a departure from previous fMRI studies investigating the neural correlates of uncertainty or confidence in perception ^64^ and decision-making ^65,66^. These previous studies used linear regression to identify brain regions whose activity increased (or decreased) with uncertainty report or confidence rating, thereby identifying the brain regions that represent uncertainty by a ‘rate code’ (i.e., increasing or decreasing averaged response amplitude with uncertainty or confidence). Perhaps such regions act as a downstream decoder and extract the uncertainty information represented by the neural populations that encode the stimulus features, in a way similar to how we decode uncertainty from voxel activity. How the regions with different coding schemes for uncertainty—the probabilistic population code reported here and the rate code described in these previous studies—interact is still an open question.

The decodable VWM signals across nearly all retinotopic ROIs in both experiments are generally consistent with the notion that the storage of VWM content involves a widespread cortical network ^24,27,50,67^. Nonetheless, the quality of VWM representations varies greatly across different brain regions. In dorsal extrastriate cortex, V3AB and its neighboring regions IPS0 and V3 showed the highest performance in decoding the memorized target locations. The standard deviations of decoding error distributions were about one third of that of the region with the lowest decoding performance. Moreover, decoding error and decoded uncertainty from V3AB and IPS0 exhibited the strongest correlations with behavioral memory error and uncertainty judgement respectively. Thus, these regions could be most critical for maintaining the content of spatial VWM. These results converge with recent studies on mental imagery and episodic memory: Breedlove et al. ^68^ built an encoding model to predict the brain activity corresponding to different ‘imagined’ images. They found higher prediction accuracy in higher-level visual areas (V3AB and IPS) than in early visual cortex. Similarly, Favila et al. ^69^ found that during retrieval of spatial positions from episodic memory, the spatially-localized memory-evoked responses in extrastriate regions V3AB and hV4 were more precise than those observed in early visual cortex. Together, their and our results highlight the importance of dorsal mid- and high-level visual cortex in maintaining behaviorally relevant information in the absence of bottom-up inputs.

Despite the centrality of the prefrontal cortex to working memory theory ^70,71^, the inferior and superior branches of the precentral sulcus (iPCS and sPCS) had the lowest decoding performance, and only in sPCS did decoding error correlate with memory error. In addition, across participants the decoding quality from frontal cortex did not predict how well a participant performed in the VWM tasks (**Fig. 8** and **Supplementary Fig. 8**). We choose sPCS and iPCS as ROIs because in the frontal cortex they exhibit the clearest retinotopic organization ^48^, and the strongest decodable spatial VWM signals in previous fMRI studies ^16,23,25^. The sPCS is believed to be the human homologue of monkey frontal eye field (FEF) ^72,73^, a macaque region known to be critically involved in spatial VWM ^74–76^, covert attention orienting ^77,78^ and saccadic eye movement ^79,80^. Perhaps surprisingly, our results indicate that compared to dorsal high-level visual cortex and IPS0, sPCS contained a quite coarse representation of memorized locations.

In a previous study, van Bergen et al. ^42^ developed and applied the Bayesian decoding method used here to quantify sensory uncertainty from early visual cortex activation patterns (voxels pooled across V1, V2 and V3) evoked by visual stimuli. They found that the uncertainty decoded from early visual cortex correlated with participants’ behavioral variability and errors in an orientation estimation task. Here, we did not observe a correlation between the decoded uncertainty and the variability (or error) of memory reports within individuals (when the individual-participant means were removed) (**Supplementary Fig. 7**). This discrepancy might reflect differences in how locations and orientations are encoded. Perhaps the correlation between decoded uncertainty and behavioral variability reported by van Bergen et al ^42^ indicates that their observers did not solely use the posterior mean when reporting orientation. For example, the use of a “posterior probability matching” strategy (reporting an orientation by drawing a sample from the posterior distribution e.g., ^81^) would increase the correlation between uncertainty and behavioral variability. Uncertainty and error (or bias) would correlate if an observer weighted the prior information more when the uncertainty was high. It is well documented that in orientation estimation, observers employ a prior reflecting the statistics of the orientations in the natural environment (e.g., more cardinal than oblique orientations ^82,83^), which is different from the statistics of the orientations used in van Bergen et al. ^42^ (e.g., uniform distribution). In the case of spatial VWM employed here, it is unlikely that observers utilized a prior for encoding locations and instead assumed that objects appeared uniformly at all possible locations (polar angles).

In a number of cortical areas, we observed strong correlations between participants’ average decoded uncertainty (across all trials) and their average memory error (across all trials; **Fig. 8**). Participants who on average represented remembered locations more precisely in their neural activation patterns—that is, with lower decoded uncertainty—were those whose working memory was more precise. This result was consistent across both experiments, and the result stands when we used an alternative index, the standard deviation of the distribution of memory errors, to quantify memory precision (**Supplementary Fig. 8**). These findings support previous studies that identified cross-subject correlations between average decoding performance and average behavioral performance ^19,22,29^. Overall, the strong cross-subject correlations we observed demonstrated that our model-based decoding approach not only provided unprecedented accuracy of decoding single-trial spatial VWM content, but also extracted features of individuals’ neural circuitry that constrained individual working memory performance. Working memory abilities predict a number of cognitive and intellectual functions, suggesting that it might be a core component upon which many high-level cognitive abilities depend ^84,85^. Although the neural sources of these individual differences in working memory remain elusive, our results suggest that the noise in the population-encoding may be an important neural source of the individual differences in VWM quality.

Overall, across two computational neuroimaging experiments we demonstrated that humans encode working memory representations as a probability distribution maintained via the activity patterns in posterior parietal and extrastriate visual cortex. These results extend previous studies identifying probabilistic sensory representations during perceptual processing and establish that probabilistic population codes are an efficient and general neural coding principle used to support higher cognitive behaviors like working memory.

## Methods

### Participants

Thirteen participants took part in Experiment 1 (two authors). Data of two participants were excluded because the eye tracking data were too noisy for extracting gaze positions reliably. Nine participants from Experiment 1 and five additional participants joined in Experiment 2. All participants had normal or corrected-to-normal vision. The experiments were conducted with the written, informed consent of each participant. The experimental protocols were approved by the University Committee on Activities involving Human Subjects at New York University, and participants received monetary compensation ($30/hr).

### Procedures

#### Experiment 1

Participants performed a memory-guided saccade task in the fMRI scanner. Each trial started with the onset of the working memory target (light gray dot) with a duration of 500 ms followed by a delay period of 12000 ms. Participants were required to remember the location of the target, and hold their gaze at the fixation point at the screen center until the end of the delay period. After the delay period, the response cue, the fixation point changing from a light gray circle outline into a filled light gray circle, instructing participants to make a saccadic eye movement to the remembered location. 700 ms after the onset of the response cue, a feedback stimulus (a white dot) was presented at the target location for 800 ms. Participants made a saccade to the feedback dot before moving their eyes back to the screen center. The intertrial interval was pseudo-randomly chosen from a range between 6, 9, or 12 seconds. Each participant completed 304 to 496 trials (346 trials per participant on average) in 2 to 3 1.5-hr scanning sessions on separate days. Each session consisted of 9 to 10 runs, each with 16 trials evenly spanning the circular space (22.5 deg spacing). Participants were allowed to take a break between runs.

#### Experiment 2

The procedures of Experiment 2 were the same as Experiment 1 except the following: In addition to the filled black dot at the screen center, the response cue contained a dark ring of which radius matching the eccentricity of the target. Participants made a saccadic eye movement onto the ring when reporting the remembered locations. Upon the detection of the saccade offset, a dot was presented at participants’ saccade landing location. Participants held a dial in their dominant hands, and were allowed to use the dial to manually adjust the location of the dot if they felt that its location did not match the location they intended to report (e.g., due to the noisy online gaze position readout). On average, only in 14% of the trials, participants’ final reported location was the same as the location initially marked by the dot. To finalize the memory report, the participants pressed the button on a button box on the other hand. Upon the button press, an arc centered at the dot (reported location) appeared on the ring. In a post-estimation wager, the participants used the dial to adjust the length of the arc. Spinning the dial clockwise increased the length of the arc along the ring and vice versa. Participants were instructed to reflect the uncertainty of their memory on the length of the arc, the longer the arc, the more uncertain. Participants finalized the arc length by pressing a button and then the feedback stimulus (a white dot) appeared at the true target location. The number of points earned by participants for each trial was displayed on the screen along with the feedback stimulus. Participants were rewarded with some points only if the true target location fell within the arc. The number of points was 100*e*^−0.08*d*^, in which *d* was the length of the arc in polar angle (°). That is, the number of points they could gain decreased exponentially with the length of the arc. To gain more points, an optimal observer should increase the length of the arc with their uncertainty. Participants were well-informed regarding the structure of the betting game and the policy of reward. Each participant completed 180 to 270 trials (227 trials per participant on average) in 2 to 3 2-hr scanning sessions on separate days.

#### Passive viewing control experiment

We scanned a subset of participants (n = 3) on an additional control experiment in which we presented a high-contrast, salient flickering checkerboard stimulus at the same locations as the WM target stimuli while participants performed a demanding discrimination task at fixation. Trial timing was identical to that used in Experiments 1 and 2. Instead of a dim target stimulus, we presented a full-contrast flickering checkerboard (0.875 deg radius; 1 cycle/deg spatial frequency; 8 Hz flicker) for 500 ms, followed by a 12 s ‘delay’ period. Throughout the trial, including during the stimulus presentation period, participants carefully attended a rapidly-flashing “+” stimulus at fixation (4 Hz) to detect targets defined by a widening or heightening of the “+” and responding with one button for each target type. We adjusted the aspect ratio of the fixation discrimination stimulus across scanning runs to maintain performance ~75%. At the end of the 12 s “delay” period, the fixation task concluded and participants received feedback about their detection performance via green/red/yellow dots (for correct/incorrect/missed responses) presented around fixation. Each of the 3 participants performed 2 sessions of this task, totaling 20-24 runs per participant.

### Setup and stimuli

Visual stimuli were presented by an LCD (VPixx ProPix) projector located behind the scanner bore, and were viewed by participants through an angled mirror with a field of view of 52° by 31°. A gray circular aperture with a diameter of 30° was presented on the screen throughout the experiments. The working memory target was a light gray dot with a diameter of 0.65°. It had an eccentricity at 12° from the central fixation point and its polar angle was pseudo-randomly chosen from 1 of 16 locations that evenly tiled the full circle within each run, and the polar angle offset of the 16 locations was varied across runs.

### Eyetracking

For all imaging sessions, we measured eye position using an EyeLink 1000 Plus infrared video-based eye tracker (SR Research) mounted beneath the screen inside the scanner bore operating at 500 Hz. The camera always tracked the participant’s right eye, and we calibrated using either a 13-point (Experiment 1, Experiment 2, and the passive viewing experiment) or 5-point (retinotopy) calibration routine at the beginning of the session and as necessary between runs. We monitored gaze data and adjusted pupil/corneal reflection detection parameters as necessary during and/or between each run.

### Behavioral data analysis

For Experiment 1, we used gaze position estimated from eye position traces as our measurement of VWM performance. We preprocessed raw gaze data using fully-automated procedures implemented within iEye_ts (github.com/tommysprague/iEye_ts) to remove blinks, adjust for drift over the course of a run, recalibrate gaze data trial-by-trial, automatically identify memory-guided saccades, and flag trials for rejection (for behavioral analyses).

We defined blinks as 200 ms before and after periods when pupil size fell below the 1.5th percentile of the distribution across all pupil size samples of the entire run (396 s). We computed velocity based on smoothed gaze time courses (5 ms standard deviation Gaussian kernel). We defined saccades based on a velocity threshold of 30 deg/s and a minimum duration of 0.0075 s and 0.25 deg amplitude. We defined periods between saccades as fixations. We drift-corrected each trial based on the modal fixation position during the trial period before the go cue appeared. To recalibrate gaze traces on each trial, we found the nearest fixation to the known target position during the feedback period (800 ms during which target was re-presented and participants were instructed to fixate this position) and fit a 3rd-order polynomial for each coordinate (X,Y) to map between actual WM position and measured gaze coordinate. We used this polynomial to recalibrate the X and Y traces across all trials within each run. We used trials for which measured gaze position was within 2.5 deg visual angle of the feedback target location for fitting the polynomial, but all trials were subjected to the resulting recalibration.

We quantified WM error based on the endpoint of the large saccadic eye movement towards the remembered position (> 5 deg amplitude, < 150 ms saccade duration), which we call the ‘primary saccade’, and the final eye position before the feedback stimulus appeared (‘final saccade’). On trials in which a subsequent corrective saccade is not made before the feedback stimulus appeared, these positions were considered identical. For both primary and final saccades, the saccade must have both begun and ended during the response period. Moreover, we exclude trials in which participants initiate a saccade faster than 100 ms after the response cue appeared ^86^.

We flagged trials for exclusion based on: (1) failures of automatic drift correction and/or excessive necessary drift correction (beyond 2.5 deg), (2) fixation outside a 2.5 deg aperture around fixation during the delay period, (3) ill-defined primary saccade, or (4) excessive error for primary saccade (> 5 degree visual angle). We included all trials for fMRI data analyses regardless of behavioral exclusion criteria during model estimation to ensure a balanced sampling of spatial positions, but only included trials with reliable behavioral estimates for all subsequent analyses including quantifying the decoding performance and correlating decoded results with behaviors (Fig. 1–8).

The analysis for Experiment 2 was similar to that of Experiment 1, with a few exceptions. As participants were allowed to use the dial to manually adjust the remembered location, we used the final dot location after the manual adjustment as the participants’ memory reports. Different from the definition of excessive error (> 5 degree visual angle) used in Experiment 1, we computed the memory error and the reported arc length in unit of degree polar angle and excluded the trials with memory error exceeding the mean error plus three standard deviations. The same exclusion criterion was applied to the trials with excessive reported arc length.

For both experiments, when quantifying participants’ behavioral memory error (Fig. 1, 4, 5 and 7), we computed the error as the (signed) difference between the reported location and WM target position in polar angle.

### Retinotopic mapping and the identification of region of interest (ROI)

Each participant was scanned for one 1.5-2 hour fMRI session for retinotopic mapping. The experimental procedures followed those reported by Mackey et al. ^48^. Participants maintained fixation at the screen center while covertly tracking a bar aperture sweeping across the screen in discrete steps and in four directions: a vertical aperture moving from the left to the right, or from the right to the left of the screen; a horizontal aperture moving from the top to the bottom, or from the bottom to the top of the screen. The bar aperture was divided into three rectangular segments (defined as a central segment and two flanking segments) with equal sizes, each containing a random dot kinematogram (RDK). Participants’ task was to discriminate in which one of the two flanking segments, the motion direction of the RDK was in the same direction as the one within the central segment. The dot motions of all the three segments changed with each discrete step. Participants reported their answer by a button press before the bar moved into the next step. The coherence of the random dot motion was staircased in order to keep the difficulty of the task at about 75% accuracy. Each session contained eight to nine runs. In each run, the bar aperture swept across the screen 12 times, and each swept consisted of 12 discrete steps. The four sweeping directions were interleaved and randomized within each run. While Mackey et al ^48^ presented different bar widths in different scanning runs, here we interleaved 3 different bar widths during the same run.

We fit a population receptive field (pRF) model with compressive spatial summation to the BOLD time series of the retinotopic mapping data for each participant ^49,87^ after smoothing on the surface (5 mm FWHM Gaussian kernel). We visualized on the cortical surface the voxels’ preferred phase angle and eccentricity estimated by the pRF model. To define the ROIs, we set a threshold to only include voxels with greater than 10% response variability explained by the pRF model. We then drew ROIs by visual inspection, primarily by identifying reversals of the voxels’ preferred phase angle on the cortical surface. We define bilateral dorsal visual ROIs V1, V2, V3, V3AB, IPS0, IPS1, IPS2, IPS3, iPCS and sPCS, each with a full visual field representation.

### MRI acquisition

MRI data were acquired on a Siemens Prisma 3T scanner with a 64-channel head/neck coil. We collected functional imaging for the working memory experiments and the passive viewing experiment with 44 slices and a voxel size of 2.5^3^ mm (4x simultaneous-multi-slice acceleration; FoV 200 × 200 mm, no in-plane acceleration, TE/TR: 30/750 ms, flip angle: 50 deg, Bandwidth: 2290 Hz/pixel; 0.56 ms echo spacing; P→ A phase encoding). Intermittently throughout each scanning session we acquired pairs of spin-echo images in the forward and reverse phase-encoding direction with identical slice prescription and no simultaneous-multi-slice acceleration (TE/TR: 45.6/3537 ms; 3 volumes per phase encode direction). These pairs are used to estimate a field map used to correct for local spatial distortions. The slice prescription was approximately parallel to the calcarine sulcus and covered most of the occipital lobe and the parietal lobe, with the exception of ventral temporal poles and ventral orbitofrontal cortex in some participants. The functional imaging data for retinotopic mapping was acquired in a separate session at a higher resolution, with a slice prescription spanning 56 slices (4x simultaneous multislice acceleration) and a voxel size of 2^3^ mm (FoV 208 x 208 mm, no in-plane acceleration, TE/TR: 36/1200 ms, flip angle: 66 deg, Bandwidth: 2604 Hz/pixel (0.51 ms echo spacing), P→A phase encoding).

For each participant, in the retinotopic mapping session, we also collected 2 or 3 T1 weighted whole-brain anatomical scans (MPRAGE sequence; 0.8 mm^3^).

### MRI data preprocessing

T1-weighted anatomical images were segmented and cortical surfaces were constructed using Freesurfer (v6.0). Functional data (EPI time series) of both the retinotopic mapping experiment and the VWM experiments were preprocessed by customized scripts using functions provided by AFNI. We applied B0 field map correction and reverse-polarity phase-encoding (reverse blip) correction to the functional data. Spatial smoothing (5 mm FWHM on the cortical surface) was only applied to the retinotopic mapping data. All the functional data were motion-corrected (6-parameter affine transform), aligned to the anatomical images, projected onto the cortical surface, then re-projected into volume space. This process incurs a small amount of smoothing along vectors perpendicular to the cortical surface, but no additional smoothing was applied. When possible, all linear and nonlinear spatial transformations were concatenated into a single transform operation to minimize additional smoothing. Linear trends were removed from the time series. For the VWM experiments, the time series of each voxel was first converted into percentage signal change for each run, and then normalized (z-score) across time points within each run.

### Generative model

We decoded the content of WM using a generative model proposed by van Bergen et al. ^42^ and van Bergen and Jehee ^45^. Specifically, we used the method named TAFKAP described in ^45^. Here, in this and the next section we briefly describe the critical components of the model and the model-fitting procedures. In the generative model, the multivariate voxel response given the stimulus location (polar angle) was modeled as a multivariate normal distribution. The average response (mean) of each voxel given a stimulus was determined by its tuning function (voxel response as a function of polar angle). The voxel tuning function was approximated by a weighted sum of eight basis functions that evenly tiled the location space. The basis functions are raised sinusoidal functions

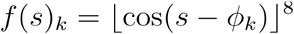

where ⌊ ⌋ represents half-wave rectification and *ϕ*_*k*_ is the center of the *k*th channel. The response of *i*th voxel *b*_*i*_ given a stimulus *s* is then modeled as

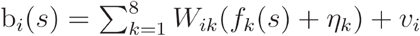

where *W* is a weighting matrix that determines the weights of each basis function for each voxel. Here, two sources of variability are considered. First, *η* is the noise specific to each basis function. This noise was carried over into each voxel by the weighting matrix **W**. It modeled the noise shared across voxels with similar voxel tuning functions. The model assumed that *η* follows a zero-mean normal distribution whose covariance matrix is a constant noise magnitude multiplied with an identity matrix *η* ~ 𝒩(0, *σ*^2^ **I**) Second, *ν* represents the noise specific to each voxel. The model assumed that the voxel-wise noise follows a zero-mean normal distribution *ν* ~ 𝒩(0, Σ). The covariance matrix of this distribution is approximated by a rank-one covariance matrix plus a diagonal matrix

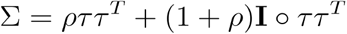

where ○ represents Hadamard product, element-wise product between two matrices. Thus, based on this generative process, the theoretical covariance matrix of the multivariate response of the voxels given a stimulus *s* is

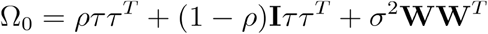

In addition to the theoretical covariance matrix, the model also considered the empirical sample covariance

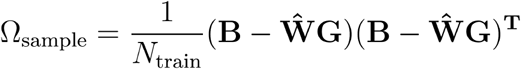

where **B** is the training data and **G** is the response of the basis functions given the training set stimuli. Thus, for each training dataset, we assumed that the voxel activity pattern followed a multivariate normal distribution.

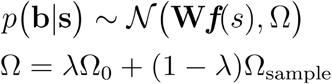

When the number of variables (voxels) is larger than the number of observations (trials), the sample covariance is not invertible. To ensure an invertible and stable estimation of the covariance matrix, here the covariance matrix was modeled as the sample covariance matrix “shrunk” ^88^ to a target covariance matrix, the theoretical covariance matrix Ω_0_. The degree of shrinkage is determined by a free parameter *λ* (see details in ^45^).

### Model fitting and decoding

For each voxel, we averaged the z-normalized percentage signal change of the BOLD time-series over a time window at 5.25 to 12.00 seconds from the delay onset. This (averaged) voxel response corresponding to the delay period was the input to the model. We trained the model and decoded spatial positions using a leave-one-run-out cross-validation procedure. For each participant, we trained the model using all the trials except those from one held-out run. We first selected the voxels with the strongest location selectivity. For each voxel, we performed a one-way ANOVA on the training dataset using the 32 target locations as a categorical independent variable and the voxel response as the dependent variable. We selected 750 voxels with the largest *F* value. These voxels were used for training the model, and later for decoding the data in the testset.

TAFKAP used a method called “bootstrap aggregating” or “bagging” to take the uncertainty of model parameters into account. Bagging is a special case of model averaging. By bootstrapping, the trials in the training dataset were resampled with replacement for multiple times to generate many bootstrap resampled datasets. Each resampled dataset had the same number of trials as the training dataset. For each resampled training dataset *j*, a set of free parameters *θ*_*j*_ was estimated by ordinary least squares. Each trial in the testset (the held-out run) was then decoded based on Bayes rule. For each trial in the testset, the posterior probability of the stimulus given the multivariate voxel response **b** was computed as

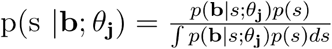

We assumed the prior *p*(*s*) to be a uniform distribution and we approximated the continuous posterior probability function by sampling 1000 steps evenly spanning the location space. The normalization factor in the posterior was computed by numerical integration. Note that for each trial in the testset, decoding was performed multiple times based on the parameters estimated using each resampled training dataset. The decoding results were averaged across all resampled training datasets to obtain one decoded posterior probability distribution

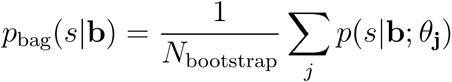

We then numerically estimated the circular mean of the posterior (*p*_bag_) to represent the decoded location, and the circular standard deviation of the posterior to represent the uncertainty of the remembered location. The number of bootstrap resampled dataset generated (*N*_bootstrap_) was determined by a stopping criterion based on Jensen-Shannon divergence (see details in ^45^).

### Statistical analysis

We tested whether the decoding error distributions (**Fig. 3A** and **Supplementary Figure 6A)** departed from a uniform distribution by a (permutation-based) V test ^89^, which is a circular variable equivalent of the Rayleigh test with the alternative hypothesis that the decoding error distributions had means centered at zero degree (polar angle). For each participant and each ROI, we computed the V statistics and compared it with the null distribution, obtained by randomly permuting the target location and then recomputing the decoding errors and their V statistics for 2000 times. The results of the V test are reported in **Supplementary Table 1** and **4**.

Decoding performance was quantified by two indices: the standard deviation of the decoding error (the decoded location minus the target location in polar angle) distribution (**Supplementary Table 2** and **5**), and the circular correlation between the decoded location and the target location (**Supplementary Table 3** and **6**). For each subject and ROI, we computed the standard deviation of the decoding error distribution and compared it with the null distribution. We obtained the null distribution by randomly permuting the target location and then recomputing the standard deviation of the decoding error for 2000 times. At the group level, we conducted the same permutation procedure to obtain the null distribution of the group-averaged standard deviation of the decoding error distribution. The same statistical procedures were applied to the circular correlation.

To relate decoding outputs to behaviors we conducted two sets of statistical tests in parallel (1) We conducted non-parametric bootstrapping to test the significance of the single-trial correlations reported in both experiments, including the circular correlation between the decoding error and memory error (**Fig. 4B** and **Fig 7B**), the correlation between decoded uncertainty and reported arc length (**Fig. 6B**), the correlation between decoded uncertainty and saccade reaction time (**Supplementary Fig. 5B and 7D**), and the correlation between decoded uncertainty and the magnitude of memory error (**Supplementary Fig. 7A and 7D**). For each ROI, we computed the correlation (or circular correlation) between the two variables in interest for each participant, and averaged the correlation coefficients across participants. We then resampled the correlation coefficients (with replacement) and computed the averaged correlation coefficients. We repeated this procedure for 2000 iterations to obtain a bootstrapped distribution of the averaged correlation coefficients. The percentage of the iterations in this distribution that was higher or lower than zero was used to compute (two-tailed) *p*-values. (2) Following the statistical tests conducted in the previous studies applying the same Bayesian decoding method ^42,45^, we also computed binned-correlation for statistical analysis (**Fig. 4C, 6C, 7C** and **Supplementary Fig. 5C, 5E, 7B, 7C, 7E and 7F**). For each participant, the trials were sorted into four bins with increasing decoding error or decoded uncertainty. The memory error or reported arc length was then computed for each bin. We then pooled data points across participants (four data points per participant) after removing the mean of each participant. For visualization, we added the grand means back to the data when plotting binned correlations. Pearson correlation coefficients were then computed based on the pooled data. We compared the correlation coefficients to the null distribution obtained by permuting the data points in the pooled data set.

We conducted permutation ANOVA to test the effect of ROI on decoding performance (**Fig. 3B** and **Supplementary Fig. 1**) and error correlations (**Fig. 4B** and **7B**). The *F*-statistic computed from the original data was compared to the null distribution of *F*-statistics, which was obtained by randomly permuting the ROI labels and calculating the *F*-statistic for 2000 times. We used a false-discovery rate (Benjamini–Hochberg procedure) for correction of multiple comparisons (the number of ROIs) with q = 0.05. We reported adjusted p-values unless otherwise specified.

## Acknowledgements

We thank New York University’s Center for Brain Imaging for technical support. This research was supported by the National Eye Institute (R01 EY-016407 and R01 EY-027925 to C.E. Curtis, F32 EY-028438 and a Sloan Research Fellowship to T. C. Sprague, and the NEI Visual Neuroscience Training Program (T32 EY007136).

## Supplementary Materials

**Supplementary Figure 1.**
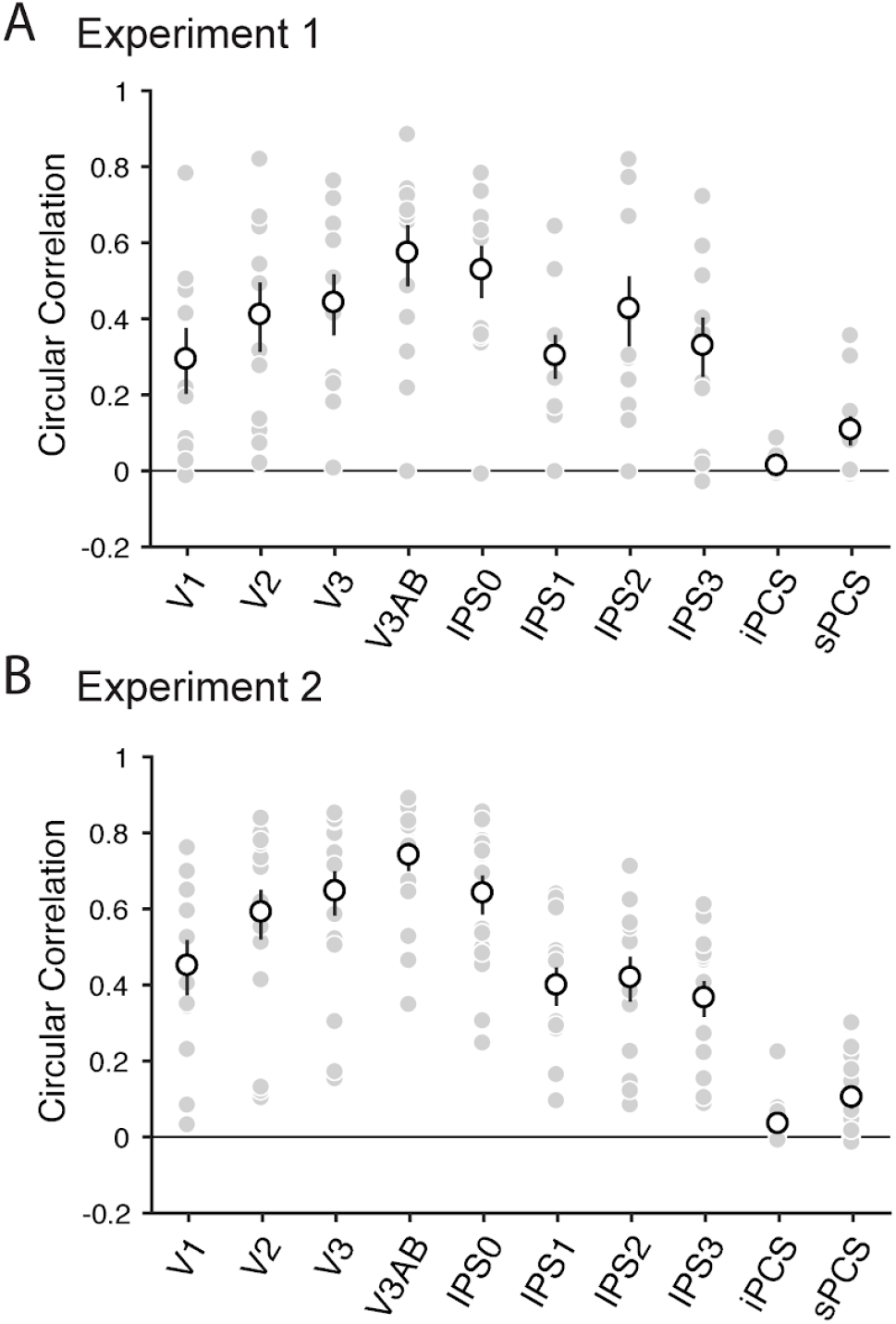
Decoded location correlates with target location. (A) Experiment 1. All ROIs showed above chance correlations (*p*<0.001 for all ROIs, permutation test). (B) Experiment 2 All ROIs showed above chance correlations (*p*<0.001 for all ROIs, permutation test). Decoding performance varied significantly across ROIs in both experiments (permutation one-way repeated-measures ANOVA, *F*(9, 90) = 17.12, *p* < 0.001, 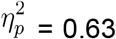 for Experiment 1; *F*(9, 117) = 48.16, *p* < 0.001, 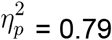 for Experiment 2).

**Supplementary Figure 2.**
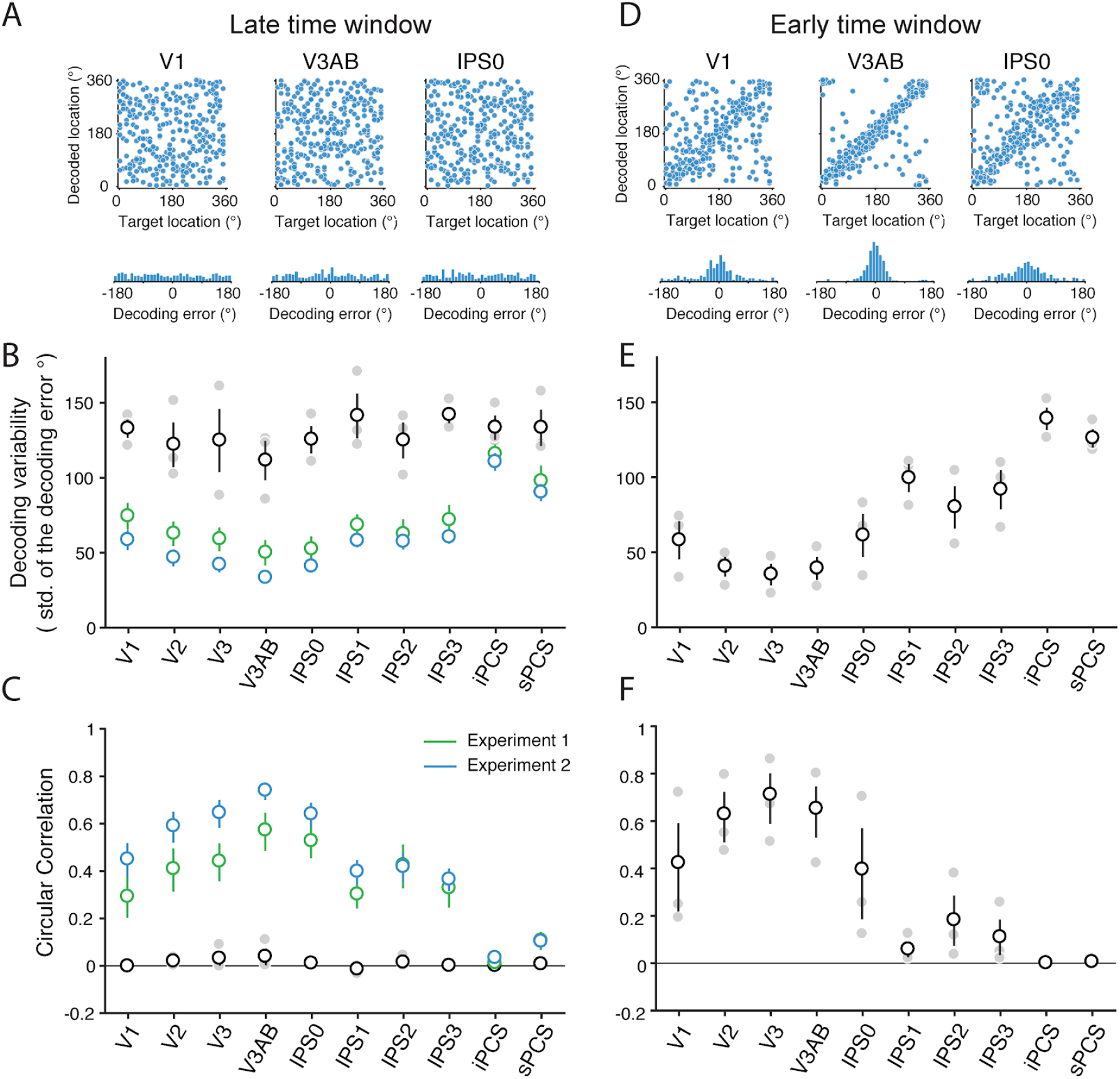
Decoding performance is low in the late time window in the passive viewing experiment. In a control experiment, a subset of participants (n = 3) performed a demanding task at fixation throughout each trial. During the same interval of the trial during which the WM target was presented in Experiment 1 and 2, we presented a high-contrast flickering checkerboard stimulus to drive strong sensory responses. (A) - (C) are the decoding results from a late time window (5.25 to 12 seconds from the delay period onset), same as the time window used for VWM analyses in Figs. 3–4 and Fig. 6–8. (A) Decoding performance of an exemplar participant, as in Fig. 3A. The top row, decoded location plotted against target location. The bottom row, the distributions of decoding error (decoded location minus the target location). (B) Decoding variability, quantified as the standard deviation of the decoding error distribution. (C) Circular correlation between the decoded location and the target location. In (B) and (C), the filled gray dots represent individual participants. The empty white dots represent group average. The error bars represent ±SEM. For comparison, the blue and the green data points represent the results of Experiment 1 and 2 using the same late time window. (D) - (F) represent the decoding results from an early time window (0.75 to 5.25 seconds from the delay period onset). Decoding performance was high for early, but not late, time periods. See Supplementary Table 1 and 2 for statistical tests on the decogin variability and circular correlation. For the late time window, decoding performance was higher in VWM experiments (Experiment 1 and 2) than in the passive viewing experiment.

**Supplementary Figure 3.**
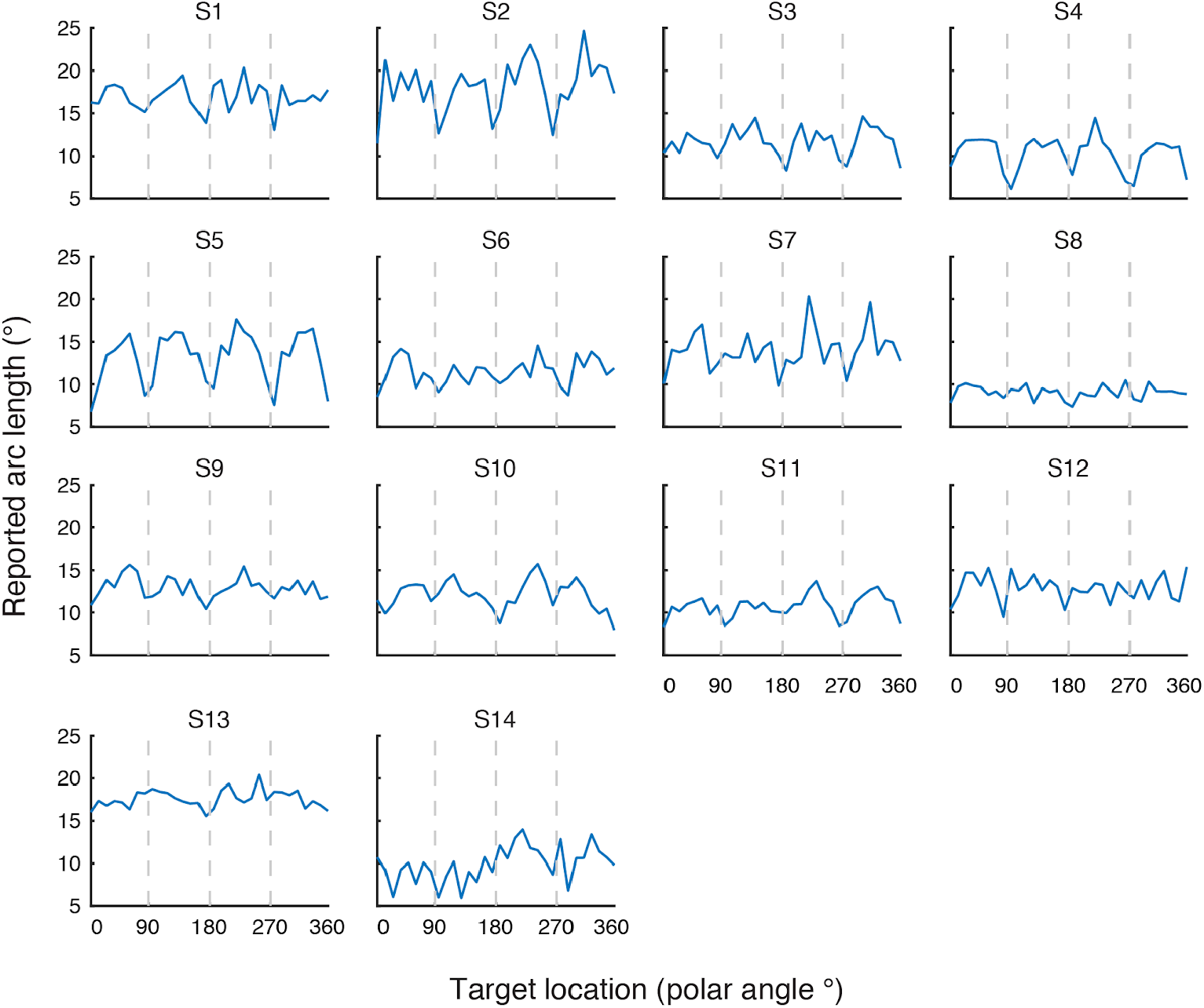
Arc length reports as a function of target location for individual participants. Here 90° corresponds to the top of the vertical meridian and 180° represents the left of the horizontal meridian.

**Supplementary Figure 4.**
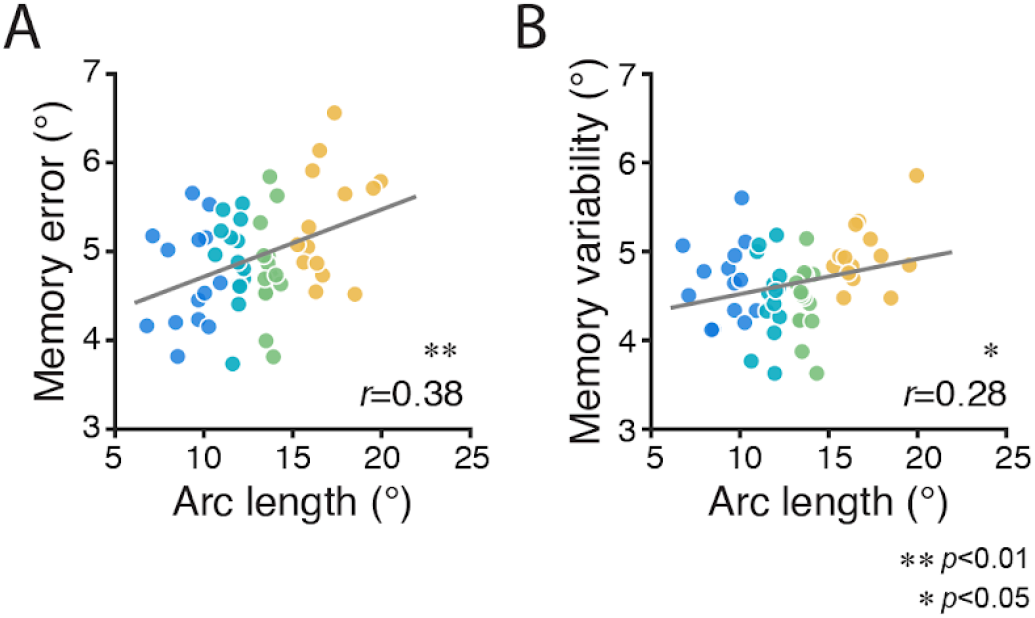
Memory error and memory variability increase with reported arc length. The data here are similar to those reported in Fig. 5D and 5E, except that the effect of target location was regressed out from the arc length. (A) Behavioral error as a function of reported arc length. Four colors represent four bins (within each of 14 participants) with increasing arc length. (B) Behavioral variability as a function of reported arc length. On trials where participants report longer arc lengths, behavioral recall of remembered positions has greater error magnitude (permutation test, *p* < 0.05) and is more variable (permutation test, *p* < 0.01).

**Supplementary Figure 5.**
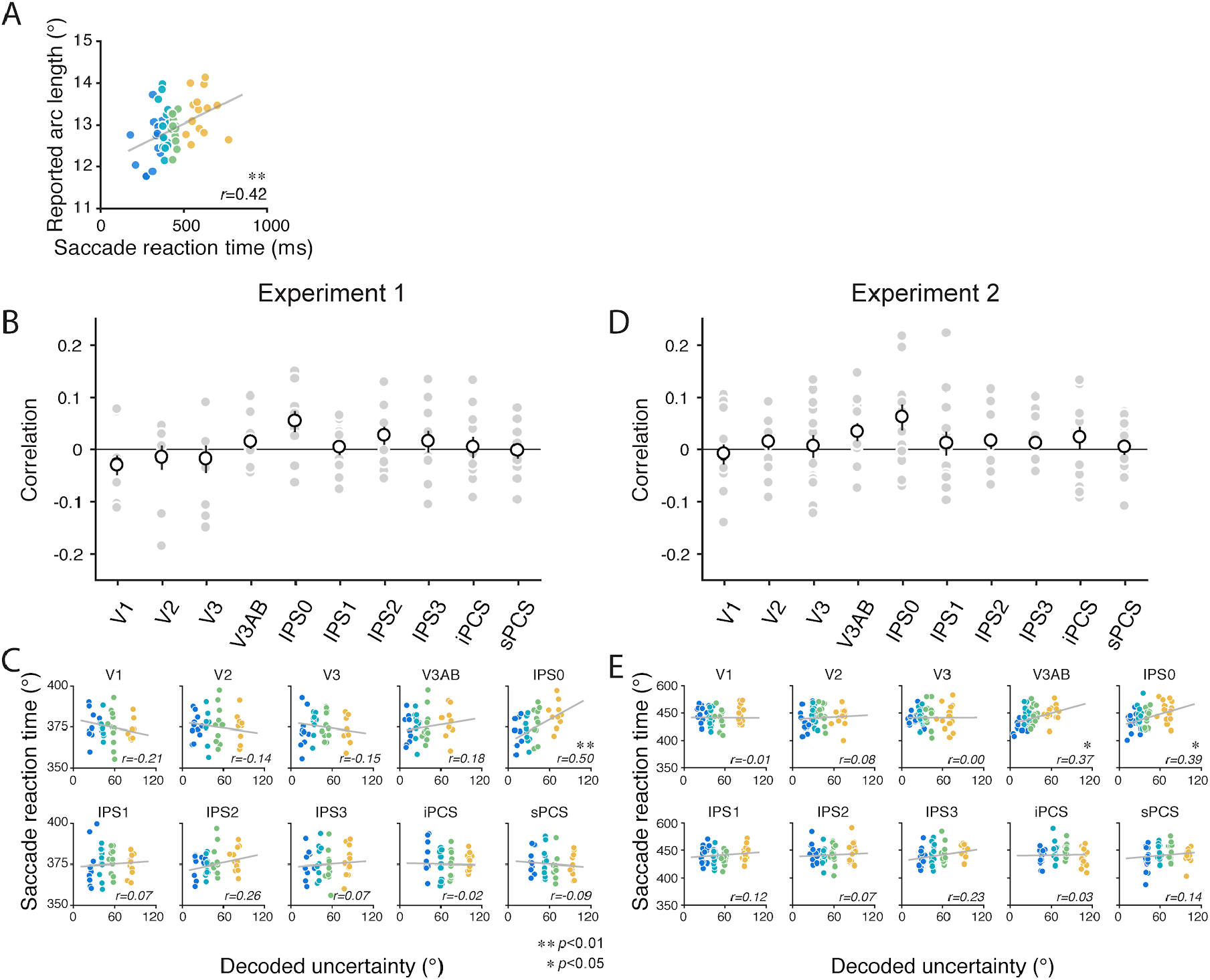
Saccade reaction time. (A) Reported arc length increased with saccade reaction time. Four colors represent four bins (within each of 14 participants) with increasing saccade reaction time. (B) Correlations between saccade reaction time and decoded uncertainty in Experiment 1. The filled gray dots represent individual participants. The empty white dots represent the group average. The error bars represent ±SEM. (C) Saccade reaction time plotted against decoded uncertainty in Experiment 1. The four colors indicate four bins (within each of 14 participants) with increasing decoded uncertainty. The gray line in each panel represents the best linear fit. The value at the lower right of each panel is the Pearson correlation coefficient. (D-E) Same analysis for Experiment 2, corresponding to (B) and (C).

**Supplementary Figure 6.**
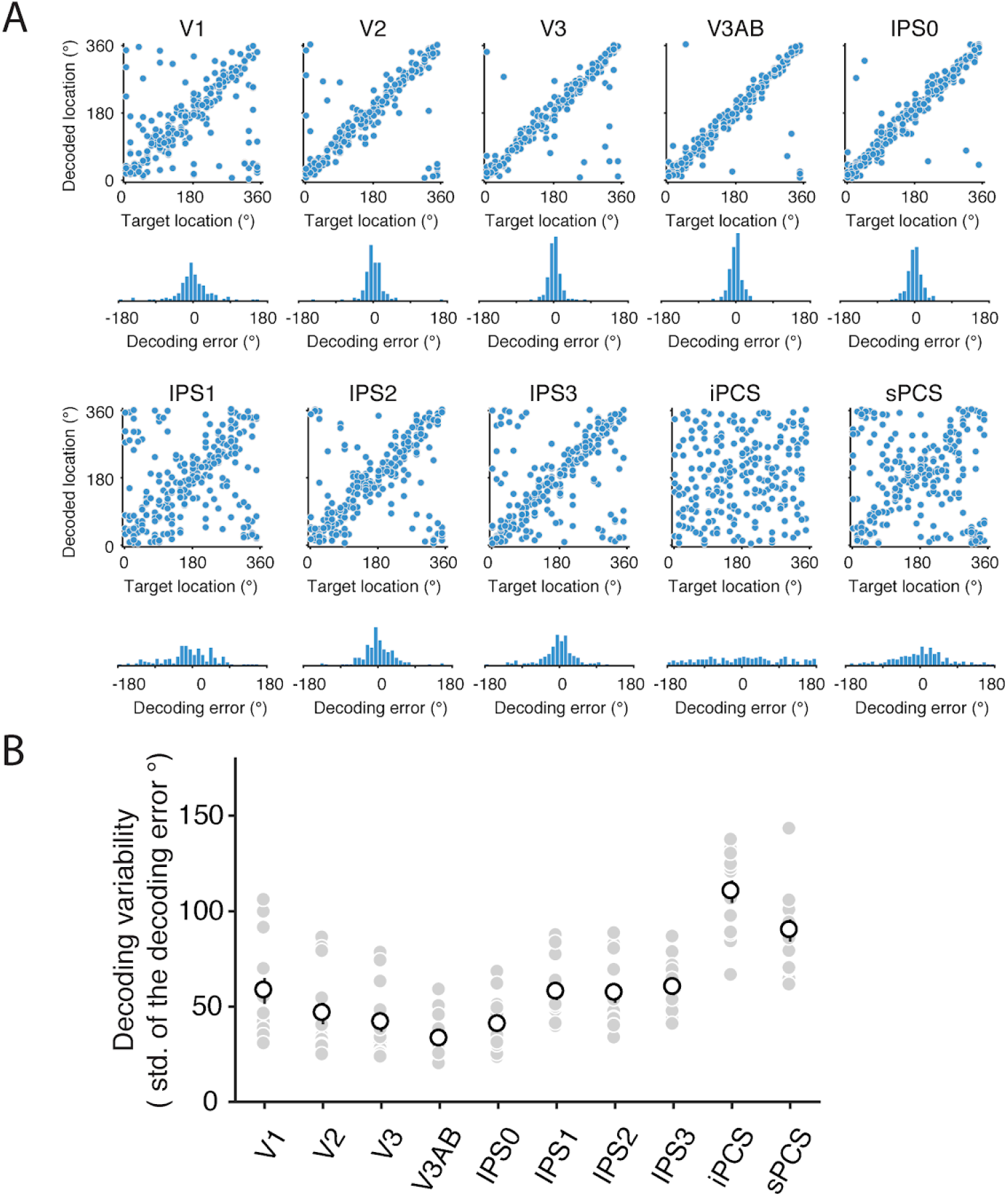
Working memory content can be precisely decoded in Experiment 2. (A) Decoding performance of an example participant. For each ROI, the top figure represents the decoded location as a function of target location. The bottom figure is the distribution of decoding error (decoded location minus the target location). (B) Decoding performance quantified as decoding variability, the standard deviation of the decoding error distribution. The filled gray dots represent individual participants. The empty white dots represent group average. The error bars represent ±SEM. Decoding performance varied significantly across ROIs (permutation one-way repeated-measures ANOVA, *p* < 0.001,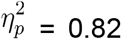).

**Supplementary Figure 7.**
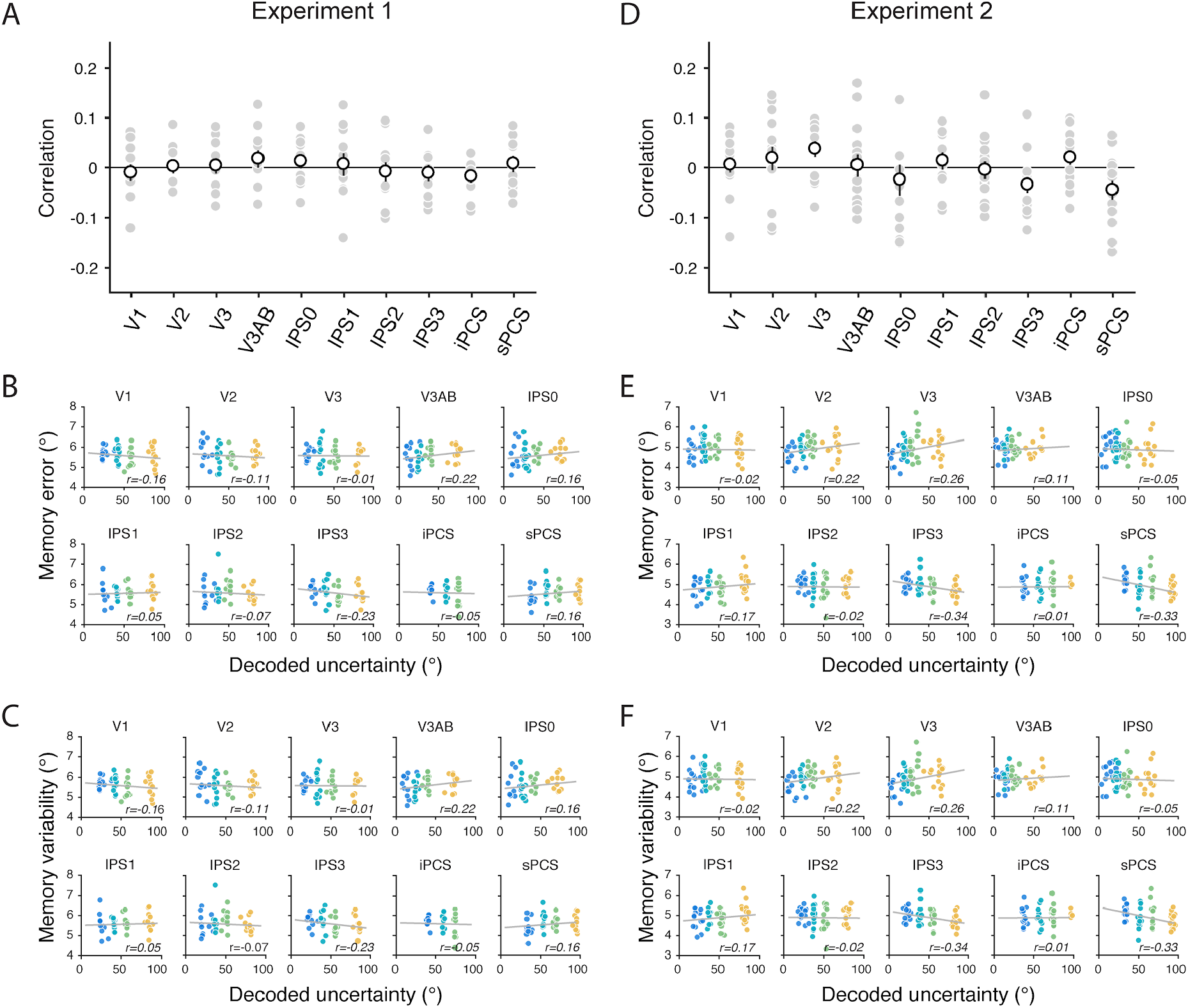
Correlations between the decoded uncertainty, memory error and memory variability. (A) Correlations are computed between decoded uncertainty and the magnitude of memory error for Experiment 1. The filled gray dots represent individual participants. The empty white dots represent the group average. The error bars represent ±SEM. (B) The magnitude of memory error plotted against decoded uncertainty. The four colors indicate four bins (within each of 14 participants) sorted by decoded uncertainty. The gray line in each panel represents the best linear fit. The value at the lower right of each panel is the Pearson correlation coefficient. (C) Similar to (B), but for each bin, the variability of memory recalls is plotted for the y-axis instead of the magnitude of error. (D-F) Correspond to (A-C) but for Experiment 2. Across (A-F) no ROI shows significant correlations between decoded uncertainty and memory error (or variability).

**Supplementary Figure 8.**
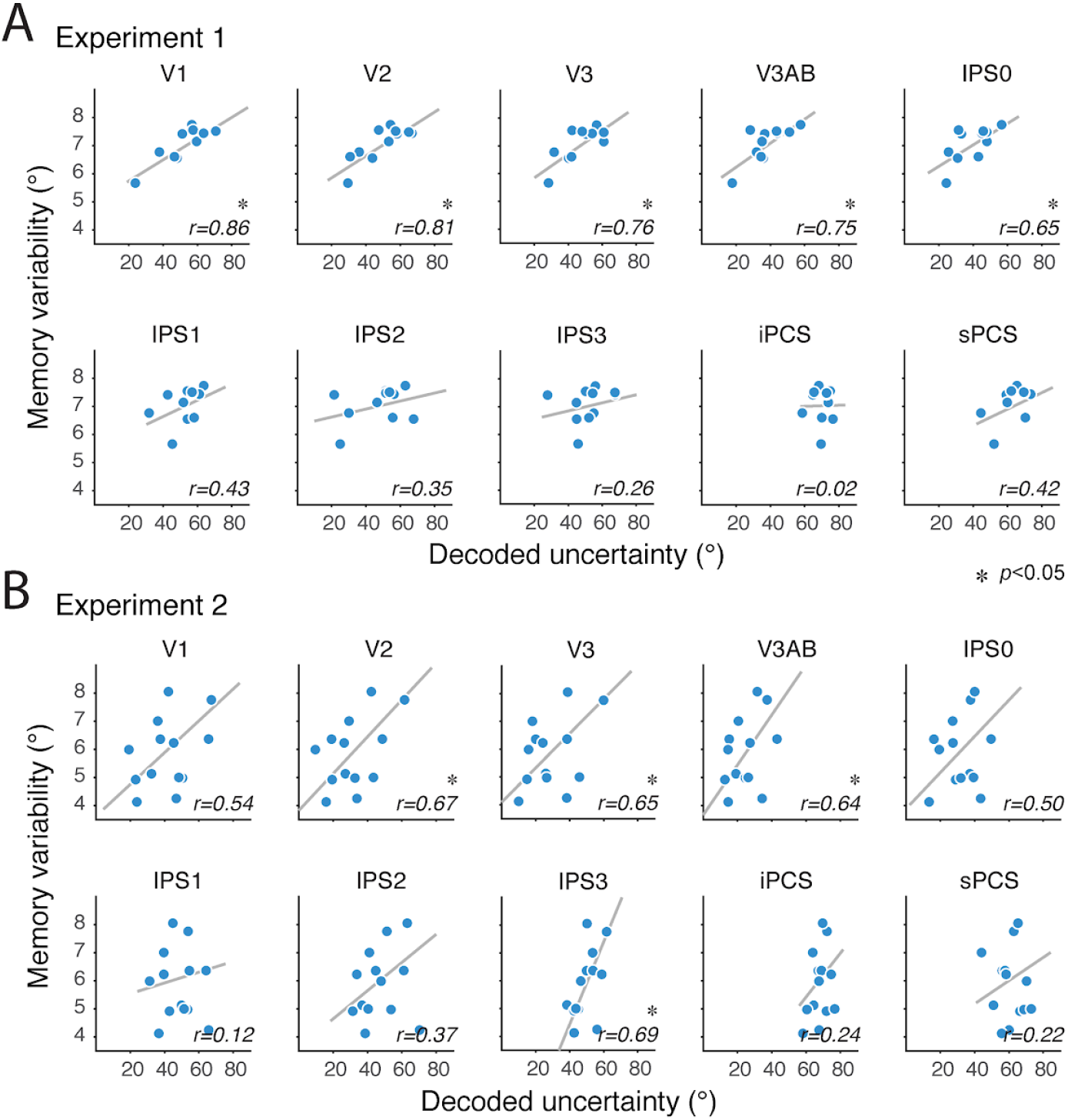
Participants with higher decoded uncertainty exhibit larger variability in memory report. . Each dot represents one participant. The decoded uncertainty (x-axis) was averaged across trials per participant. Memory variability (y-axis) is the standard deviation of behavioral error distribution of each participant. The gray lines represent the best linear fit.

**Supplementary Table 1.** Statistical test for the uniformity of the decoding error distributions in Experiment 1. For each participant and ROI, we reported a V statistics and the (uncorrected) *p*-value obtained by permutations (see Methods). A *p*-value smaller than 0.05 indicates that the error distribution is not uniform and favors the alternative hypothesis that the error distribution has a mean centered at zero degree (polar angle). The *p*-values smaller than 0.05 are highlighted by red color.

|  | V1 | V2 | V3 | V3AB | IPS0 | IPS1 | IPS2 | IPS3 | iPCS | sPCS |
| --- | --- | --- | --- | --- | --- | --- | --- | --- | --- | --- |
| S1 | 192 (<.001) | 223 (<.001) | 236 (<.001) | 240 (<.001) | 246 (<.001) | 224 (<.001) | 228 (<.001) | 199 (<.001) | 78 (<.001) | 175 (<.001) |
| S2 | 18 (0.073) | 34 (0.001) | 22 (0.022) | 19 (0.066) | 15 (0.142) | 15 (0.133) | 8 (0.408) | 8 (0.429) | -11 (0.319) | 1 (0.914) |
| S3 | 295 (<.001) | 302 (<.001) | 291 (<.001) | 314 (<.001) | 286 (<.001) | 245 (<.001) | 293 (<.001) | 256 (<.001) | 78 (<.001) | 186 (<.001) |
| S4 | 138 (<.001) | 177 (<.001) | 204 (<.001) | 256 (<.001) | 253 (<.001) | 194 (<.001) | 286 (<.001) | 269 (<.001) | 94 (<.001) | 131 (<.001) |
| S5 | 129 (<.001) | 192 (<.001) | 195 (<.001) | 224 (<.001) | 214 (<.001) | 133 (<.001) | 177 (<.001) | 164 (<.001) | 34 (0.001) | 98 (<.001) |
| S6 | 161 (<.001) | 189 (<.001) | 181 (<.001) | 254 (<.001) | 213 (<.001) | 181 (<.001) | 233 (<.001) | 211 (<.001) | 47 (<.001) | 150 (<.001) |
| S7 | 60 (<.001) | 66 (<.001) | 91 (<.001) | 119 (<.001) | 127 (<.001) | 78 (<.001) | 90 (<.001) | 40 (<.001) | 18 (0.076) | 1 (0.885) |
| S8 | 55 (<.001) | 82 (<.001) | 108 (<.001) | 142 (<.001) | 137 (<.001) | 92 (<.001) | 113 (<.001) | 107 (<.001) | 45 (<.001) | 65 (<.001) |
| S9 | 197 (<.001) | 226 (<.001) | 247 (<.001) | 261 (<.001) | 251 (<.001) | 168 (<.001) | 115 (<.001) | 194 (<.001) | 17 (0.189) | 90 (<.001) |
| S10 | 195 (<.001) | 225 (<.001) | 214 (<.001) | 228 (<.001) | 219 (<.001) | 165 (<.001) | 160 (<.001) | 136 (<.001) | 51 (<.001) | 55 (<.001) |
| S11 | 38 (0.001) | 64 (<.001) | 98 (<.001) | 109 (<.001) | 142 (<.001) | 115 (<.001) | 129 (<.001) | 34 (0.002) | 6 (0.632) | 7 (0.511) |

**Supplementary Table 2.** The variability of decoding error in Experiment 1 and the passive viewing experiment. For each participant and ROI, we report the standard deviation of the decoding error distribution (in unit of degree polar angle). The values in the parenthesis are uncorrected *p*-values computed by comparing the variability of the data with a null distribution obtained by a permutation procedure. The *p*-values smaller than 0.05 are highlighted by red color.

|  | V1 | V2 | V3 | V3AB | IPS0 | IPS1 | IPS2 | IPS3 | iPCS | sPCS |
| --- | --- | --- | --- | --- | --- | --- | --- | --- | --- | --- |
| **Experiment 1** |  |  |  |  |  |  |  |  |  |  |
| **S1** | 50 (<.001) | 38 (<.001) | 33 (<.001) | 32 (<.001) | 29 (<.001) | 38 (<.001) | 36 (<.001) | 47 (<.001) | 91 (<.001) | 55 (<.001) |
| **S2** | 127 (0.447) | 110 (0.006) | 121 (0.185) | 122 (0.222) | 131 (0.660) | 130 (0.564) | 145 (0.591) | 143 (0.692) | 140 (0.827) | 161 (0.143) |
| **S3** | 28 (<.001) | 26 (<.001) | 30 (<.001) | 20 (<.001) | 32 (<.001) | 45 (<.001) | 29 (<.001) | 42 (<.001) | 97 (<.001) | 62 (<.001) |
| **S4** | 74 (<.001) | 62 (<.001) | 54 (<.001) | 37 (<.001) | 38 (<.001) | 57 (<.001) | 26 (<.001) | 33 (<.001) | 90 (<.001) | 76 (<.001) |
| **S5** | 70 (<.001) | 48 (<.001) | 47 (<.001) | 36 (<.001) | 40 (<.001) | 69 (<.001) | 54 (<.001) | 58 (<.001) | 115 (0.024) | 82 (<.001) |
| **S6** | 73 (<.001) | 65 (<.001) | 68 (<.001) | 49 (<.001) | 60 (<.001) | 68 (<.001) | 54 (<.001) | 60 (<.001) | 116 (0.005) | 76 (<.001) |
| **S7** | 90 (<.001) | 86 (<.001) | 73 (<.001) | 60 (<.001) | 56 (<.001) | 80 (<.001) | 74 (<.001) | 103 (0.001) | 125 (0.355) | 139 (0.876) |
| **S8** | 96 (<.001) | 81 (<.001) | 69 (<.001) | 55 (<.001) | 57 (<.001) | 76 (<.001) | 67 (<.001) | 69 (<.001) | 103 (<.001) | 90 (<.001) |
| **S9** | 54 (<.001) | 45 (<.001) | 38 (<.001) | 33 (<.001) | 37 (<.001) | 63 (<.001) | 80 (<.001) | 55 (<.001) | 136 (0.688) | 90 (<.001) |
| **S10** | 47 (<.001) | 36 (<.001) | 41 (<.001) | 35 (<.001) | 39 (<.001) | 58 (<.001) | 59 (<.001) | 68 (<.001) | 105 (<.001) | 103 (<.001) |
| **S11** | 108 (0.002) | 92 (<.001) | 75 (<.001) | 71 (<.001) | 57 (<.001) | 68 (<.001) | 62 (<.001) | 112 (0.010) | 155 (0.287) | 141 (0.874) |
| **Passive viewing experiment** |  |  |  |  |  |  |  |  |  |  |
| **S3** | 142 (1.000) | 151 (0.539) | 126 (0.156) | 126 (0.143) | 142 (0.977) | 171 (0.080) | 132 (0.431) | 140 (0.894) | 149 (0.596) | 122 (0.073) |
| **S4** | 121 (0.057) | 102 (<.001) | 88 (<.001) | 85 (<.001) | 123 (0.089) | 131 (0.367) | 101 (<.001) | 133 (0.456) | 124 (0.093) | 120 (0.052) |
| **S6** | 135 (0.456) | 113 (<.001) | 161 (0.281) | 123 (0.051) | 111 (0.001) | 122 (0.041) | 141 (0.831) | 152 (0.568) | 127 (0.118) | 157 (0.349) |

**Supplementary Table 3.** Circular correlation between the decoded location and the target location in Experiment 1 and the passive viewing experiment. For each participant and ROI, we report the circular correlation and the values in the parenthesis are uncorrected *p*-values computed by comparing the correlation with a null distribution obtained by a permutation procedure. The *p*-values smaller than 0.05 are highlighted by red color.

|  | V1 | V2 | V3 | V3AB | IPS0 | IPS1 | IPS2 | IPS3 | iPCS | sPCS |
| --- | --- | --- | --- | --- | --- | --- | --- | --- | --- | --- |
| **Experiment 1** |  |  |  |  |  |  |  |  |  |  |
| **S1** | 0.47<br>($<.001$) | 0.64<br>($<.001$) | 0.71<br>($<.001$) | 0.74<br>($<.001$) | 0.78<br>($<.001$) | 0.64<br>($<.001$) | 0.67<br>($<.001$) | 0.51<br>($<.001$) | -0.01<br>(0.051) | 0.35<br>($<.001$) |
| **S2** | -0.02<br>(0.030) | 0.02<br>(0.029) | 0.00<br>(0.429) | -0.01<br>(0.324) | -0.01<br>(0.088) | -0.01<br>(0.335) | -0.01<br>(0.275) | -0.03<br>(0.001) | -0.01<br>(0.129) | -0.01<br>(0.076) |
| **S3** | 0.78<br>($<.001$) | 0.82<br>($<.001$) | 0.76<br>($<.001$) | 0.88<br>($<.001$) | 0.73<br>($<.001$) | 0.53<br>($<.001$) | 0.77<br>($<.001$) | 0.59<br>($<.001$) | 0.04<br>($<.001$) | 0.30<br>($<.001$) |
| **S4** | 0.19<br>($<.001$) | 0.31<br>($<.001$) | 0.41<br>($<.001$) | 0.65<br>($<.001$) | 0.64<br>($<.001$) | 0.36<br>($<.001$) | 0.82<br>($<.001$) | 0.72<br>($<.001$) | 0.08<br>($<.001$) | 0.11<br>($<.001$) |
| **S5** | 0.22<br>($<.001$) | 0.49<br>($<.001$) | 0.50<br>($<.001$) | 0.67<br>($<.001$) | 0.61<br>($<.001$) | 0.23<br>($<.001$) | 0.41<br>($<.001$) | 0.36<br>($<.001$) | 0.02<br>(0.010) | 0.08<br>($<.001$) |
| **S6** | 0.19<br>($<.001$) | 0.27<br>($<.001$) | 0.24<br>($<.001$) | 0.48<br>($<.001$) | 0.33<br>($<.001$) | 0.24<br>($<.001$) | 0.41<br>($<.001$) | 0.32<br>($<.001$) | 0.01<br>(0.073) | 0.15<br>($<.001$) |
| **S7** | 0.08<br>($<.001$) | 0.10<br>($<.001$) | 0.18<br>($<.001$) | 0.31<br>($<.001$) | 0.35<br>($<.001$) | 0.14<br>($<.001$) | 0.17<br>($<.001$) | 0.03<br>(0.001) | 0.01<br>(0.249) | -0.00<br>(0.754) |
| **S8** | 0.06<br>($<.001$) | 0.13<br>($<.001$) | 0.23<br>($<.001$) | 0.40<br>($<.001$) | 0.37<br>($<.001$) | 0.17<br>($<.001$) | 0.24<br>($<.001$) | 0.23<br>($<.001$) | -0.00<br>(0.476) | 0.08<br>($<.001$) |
| **S9** | 0.41<br>($<.001$) | 0.54<br>($<.001$) | 0.65<br>($<.001$) | 0.72<br>($<.001$) | 0.66<br>($<.001$) | 0.29<br>($<.001$) | 0.13<br>($<.001$) | 0.40<br>($<.001$) | -0.01<br>(0.049) | 0.08<br>($<.001$) |
| **S10** | 0.50<br>($<.001$) | 0.66<br>($<.001$) | 0.60<br>($<.001$) | 0.68<br>($<.001$) | 0.63<br>($<.001$) | 0.35<br>($<.001$) | 0.29<br>($<.001$) | 0.21<br>($<.001$) | 0.01<br>(0.039) | -0.01<br>(0.237) |
| **S11** | 0.02<br>(0.002) | 0.07<br>($<.001$) | 0.18<br>($<.001$) | 0.21<br>($<.001$) | 0.35<br>($<.001$) | 0.24<br>($<.001$) | 0.30<br>($<.001$) | 0.01<br>(0.029) | -0.00<br>(0.598) | -0.00<br>(0.697) |
| **Passive viewing experiment** |  |  |  |  |  |  |  |  |  |  |
| **S3** | -0.00 (0.603) | 0.00 (0.878) | 0.01 (0.163) | 0.00 (0.266) | 0.00 (0.971) | -0.01 (0.080) | -0.00 (0.583) | 0.00 (0.579) | -0.01 (0.054) | 0.01 (0.060) |
| **S4** | -0.01 (0.065) | 0.03 ($<.001$) | 0.09 ($<.001$) | 0.11 ($<.001$) | 0.01 (0.070) | -0.04 ($<.001$) | 0.04 ($<.001$) | -0.00 (0.833) | 0.01 (0.076) | 0.01 (0.033) |
| **S6** | 0.00 (0.422) | 0.02 ($<.001$) | -0.00 (0.237) | 0.00 (0.576) | 0.02 ($<.001$) | -0.00 (0.278) | -0.00 (0.927) | -0.00 (0.386) | -0.00 (0.779) | -0.00 (0.572) |

**Supplementary Table 4.** Statistical test for the uniformity of the decoding error distributions in Experiment 2. For each participant and ROI, we reported a V statistics and the (uncorrected) *p*-value obtained by permutations (see Methods). A *p*-value smaller than 0.05 indicates that the error distribution is not uniform and favors the alternative hypothesis that the error distribution has a mean centered at zero degree (polar angle). The *p*-values smaller than 0.05 are highlighted by red color.

|  | V1 | V2 | V3 | V3AB | IPS0 | IPS1 | IPS2 | IPS3 | iPCS | sPCS |
| --- | --- | --- | --- | --- | --- | --- | --- | --- | --- | --- |
| S1 | 56 (<.001) | 65 (<.001) | 79 (<.001) | 119 (<.001) | 99 (<.001) | 63 (<.001) | 77 (<.001) | 64 (<.001) | 35 (0.001) | 41 (<.001) |
| S2 | 158 (<.001) | 173 (<.001) | 178 (<.001) | 185 (<.001) | 182 (<.001) | 146 (<.001) | 154 (<.001) | 141 (<.001) | 70 (<.001) | 110 (<.001) |
| S3 | 176 (<.001) | 187 (<.001) | 189 (<.001) | 194 (<.001) | 154 (<.001) | 122 (<.001) | 156 (<.001) | 151 (<.001) | 23 (0.024) | 106 (<.001) |
| S4 | 157 (<.001) | 157 (<.001) | 160 (<.001) | 167 (<.001) | 155 (<.001) | 122 (<.001) | 152 (<.001) | 136 (<.001) | 43 (<.001) | 70 (<.001) |
| S5 | 170 (<.001) | 209 (<.001) | 213 (<.001) | 221 (<.001) | 216 (<.001) | 126 (<.001) | 175 (<.001) | 162 (<.001) | 26 (0.018) | 111 (<.001) |
| S6 | 51 (<.001) | 85 (<.001) | 126 (<.001) | 157 (<.001) | 129 (<.001) | 94 (<.001) | 81 (<.001) | 91 (<.001) | 13 (0.232) | 66 (<.001) |
| S7 | 39 (<.001) | 84 (<.001) | 95 (<.001) | 159 (<.001) | 161 (<.001) | 118 (<.001) | 105 (<.001) | 70 (<.001) | -16 (0.105) | 10 (0.327) |
| S8 | 104 (<.001) | 154 (<.001) | 183 (<.001) | 191 (<.001) | 181 (<.001) | 138 (<.001) | 131 (<.001) | 137 (<.001) | 68 (<.001) | 40 (<.001) |
| S9 | 104 (<.001) | 132 (<.001) | 128 (<.001) | 145 (<.001) | 120 (<.001) | 62 (<.001) | 53 (<.001) | 82 (<.001) | -10 (0.230) | 37 (<.001) |
| S10 | 199 (<.001) | 211 (<.001) | 218 (<.001) | 216 (<.001) | 218 (<.001) | 191 (<.001) | 180 (<.001) | 170 (<.001) | 123 (<.001) | 135 (<.001) |
| S11 | 119 (<.001) | 140 (<.001) | 144 (<.001) | 165 (<.001) | 138 (<.001) | 129 (<.001) | 148 (<.001) | 146 (<.001) | 12 (0.192) | 50 (<.001) |
| S12 | 132 (<.001) | 172 (<.001) | 176 (<.001) | 192 (<.001) | 192 (<.001) | 173 (<.001) | 173 (<.001) | 106 (<.001) | 12 (0.233) | 72 (<.001) |
| S13 | 140 (<.001) | 153 (<.001) | 168 (<.001) | 191 (<.001) | 166 (<.001) | 161 (<.001) | 90 (<.001) | 127 (<.001) | 18 (0.117) | 44 (<.001) |
| S14 | 181 (<.001) | 190 (<.001) | 192 (<.001) | 189 (<.001) | 180 (<.001) | 162 (<.001) | 135 (<.001) | 135 (<.001) | 63 (<.001) | 54 (<.001) |

**Supplementary Table 5.** The variability of decoding error in Experiment 2. For each participant and ROI, we report the standard deviation of the decoding error distribution (in unit of degree polar angle). The values in the parenthesis are uncorrected *p*-values computed by comparing the variability of the data with a null distribution obtained by a permutation procedure. The *p*-values smaller than 0.05 are highlighted by red color.

|  | V1 | V2 | V3 | V3AB | IPS0 | IPS1 | IPS2 | IPS3 | iPCS | sPCS |
| --- | --- | --- | --- | --- | --- | --- | --- | --- | --- | --- |
| S1 | 91 (<.001) | 86 (<.001) | 78 (<.001) | 58 (<.001) | 68 (<.001) | 87 (<.001) | 79 (<.001) | 86 (<.001) | 106 (0.002) | 102 (<.001) |
| S2 | 41 (<.001) | 34 (<.001) | 31 (<.001) | 26 (<.001) | 29 (<.001) | 48 (<.001) | 43 (<.001) | 50 (<.001) | 84 (<.001) | 64 (<.001) |
| S3 | 38 (<.001) | 32 (<.001) | 31 (<.001) | 28 (<.001) | 48 (<.001) | 62 (<.001) | 47 (<.001) | 49 (<.001) | 121 (0.167) | 69 (<.001) |
| S4 | 30 (<.001) | 30 (<.001) | 27 (<.001) | 22 (<.001) | 31 (<.001) | 50 (<.001) | 33 (<.001) | 43 (<.001) | 97 (0.001) | 79 (<.001) |
| S5 | 46 (<.001) | 27 (<.001) | 25 (<.001) | 20 (<.001) | 23 (<.001) | 64 (<.001) | 44 (<.001) | 49 (<.001) | 120 (0.114) | 70 (<.001) |
| S6 | 99 (0.001) | 81 (<.001) | 63 (<.001) | 50 (<.001) | 62 (<.001) | 77 (<.001) | 82 (<.001) | 78 (<.001) | 128 (0.426) | 89 (<.001) |
| S7 | 105 (0.002) | 79 (<.001) | 74 (<.001) | 45 (<.001) | 44 (<.001) | 63 (<.001) | 69 (<.001) | 86 (<.001) | 122 (0.198) | 143 (0.714) |
| S8 | 69 (<.001) | 47 (<.001) | 33 (<.001) | 29 (<.001) | 34 (<.001) | 55 (<.001) | 57 (<.001) | 55 (<.001) | 87 (<.001) | 105 (0.002) |
| S9 | 59 (<.001) | 44 (<.001) | 46 (<.001) | 36 (<.001) | 50 (<.001) | 83 (<.001) | 88 (<.001) | 71 (<.001) | 132 (0.808) | 100 (0.001) |
| S10 | 35 (<.001) | 29 (<.001) | 25 (<.001) | 26 (<.001) | 25 (<.001) | 38 (<.001) | 43 (<.001) | 47 (<.001) | 66 (<.001) | 61 (<.001) |
| S11 | 55 (<.001) | 44 (<.001) | 42 (<.001) | 29 (<.001) | 45 (<.001) | 49 (<.001) | 40 (<.001) | 40 (<.001) | 124 (0.350) | 93 (<.001) |
| S12 | 58 (<.001) | 40 (<.001) | 38 (<.001) | 30 (<.001) | 29 (<.001) | 39 (<.001) | 40 (<.001) | 69 (<.001) | 137 (0.995) | 85 (<.001) |
| S13 | 59 (<.001) | 54 (<.001) | 48 (<.001) | 38 (<.001) | 49 (<.001) | 51 (<.001) | 80 (<.001) | 64 (<.001) | 130 (0.497) | 105 (0.001) |
| S14 | 30 (<.001) | 24 (<.001) | 23 (<.001) | 25 (<.001) | 31 (<.001) | 41 (<.001) | 53 (<.001) | 53 (<.001) | 88 (<.001) | 94 (<.001) |

**Supplementary Table 6.**
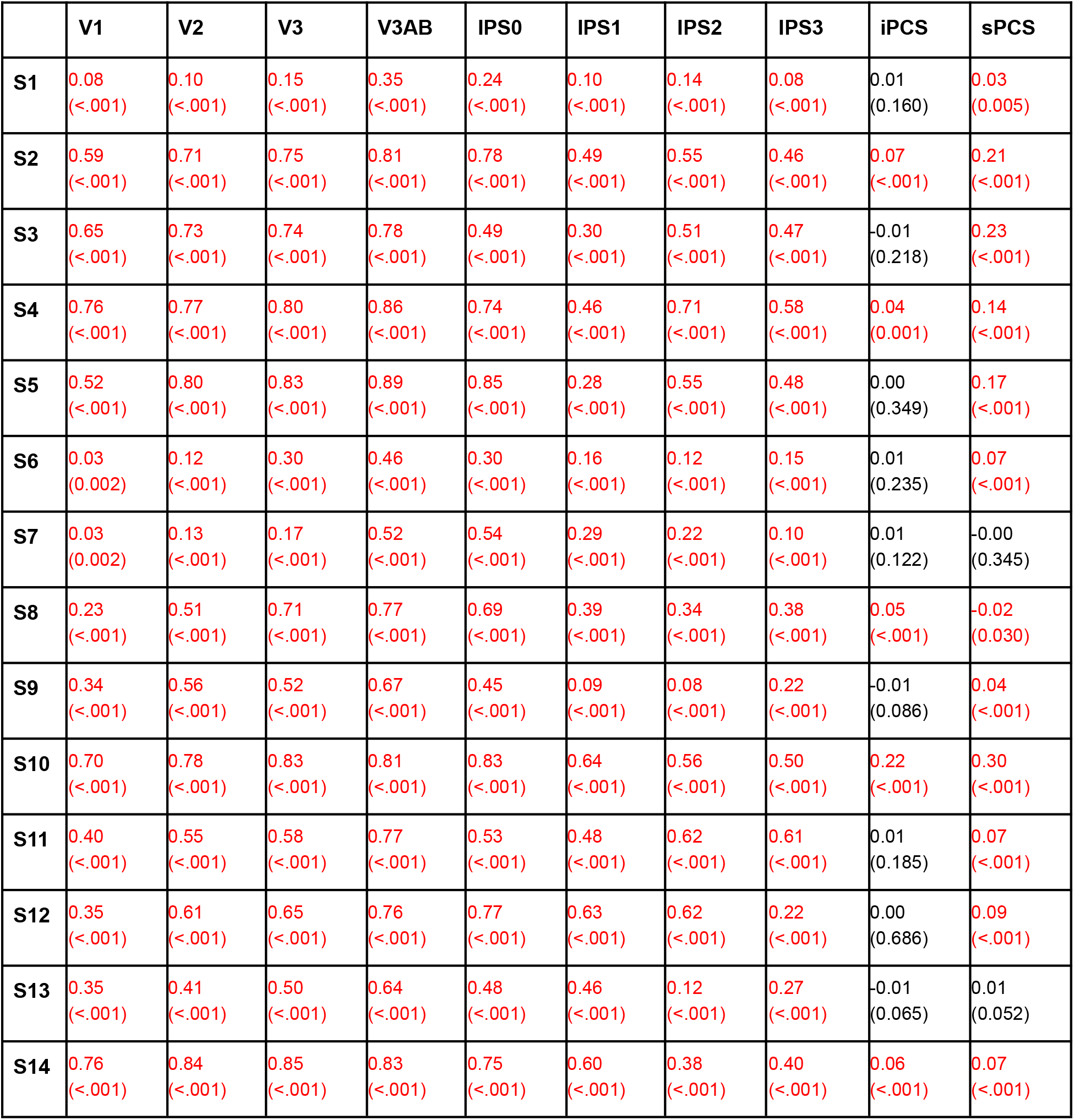
Circular correlation between the decoded location and the target location in Experiment 2. For each participant and ROI, we report the circular correlation and the values in the parenthesis are uncorrected p-values computed by comparing the correlation with a null distribution obtained by a permutation procedure. The *p*-values smaller than 0.05 are highlighted by red color.

## References

1. Wagner, A. D. Working memory contributions to human learning and remembering. Neuron 22, 19–22 (1999).

2. Collins, A. G. E. & Frank, M. J. How much of reinforcement learning is working memory, not reinforcement learning? A behavioral, computational, and neurogenetic analysis. Eur. J. Neurosci. 35, 1024–1035 (2012).

3. Curtis, C. E. & Lee, D. Beyond working memory: the role of persistent activity in decision making. Trends Cogn. Sci. 14, 216–222 (2010).

4. Ma, W. J., Husain, M. & Bays, P. M. Changing concepts of working memory. Nat. Neurosci. 17, 347–356 (2014).

5. Fougnie, D., Suchow, J. W. & Alvarez, G. Variability in the quality of visual working memory. Nature Communications 3, 1229 (2012).

6. Keshvari, S., van den Berg, R. & Ma, W. J. Probabilistic computation in human perception under variability in encoding precision. PLoS One 7, e40216 (2012).

7. Rademaker, R. L., Tredway, C. H. & Tong, F. Introspective judgments predict the precision and likelihood of successful maintenance of visual working memory. J. Vis. 12, 21 (2012).

8. van den Berg, R., Yoo, A. H. & Ma, W. J. Fechner’s law in metacognition: A quantitative model of visual working memory confidence. Psychol. Rev. 124, 197–214 (2017).

9. Samaha, J. & Postle, B. R. Correlated individual differences suggest a common mechanism underlying metacognition in visual perception and visual short-term memory. Proc. Biol. Sci. 284, (2017).

10. Devkar, D., Wright, A. A. & Ma, W. J. Monkeys and humans take local uncertainty into account when localizing a change. J. Vis. 17, 4 (2017).

11. Yoo, A. H., Acerbi, L. & Ma, W. ji. Uncertainty is Maintained and Used in Working Memory. Cold Spring Harbor Laboratory 2020.10.06.328310 (2020) doi:10.1101/2020.10.06.328310.

12. Yoo, A. H., Klyszejko, Z., Curtis, C. E. & Ma, W. J. Strategic allocation of working memory resource. Sci. Rep. 8, 16162 (2018).

13. Honig, M., Ma, W. J. & Fougnie, D. Humans incorporate trial-to-trial working memory uncertainty into rewarded decisions. Proc. Natl. Acad. Sci. U. S. A. 117, 8391–8397 (2020).

14. Serences, J. T., Ester, E. F., Vogel, E. K. & Awh, E. Stimulus-specific delay activity in human primary visual cortex. Psychol. Sci. 20, 207–214 (2009).

15. Harrison, S. A. & Tong, F. Decoding reveals the contents of visual working memory in early visual areas. Nature 458, 632–635 (2009).

16. Jerde, T. A., Merriam, E. P., Riggall, A. C., Hedges, J. H. & Curtis, C. E. Prioritized maps of space in human frontoparietal cortex. J. Neurosci. 32, 17382–17390 (2012).

17. Riggall, A. C. & Postle, B. R. The relationship between working memory storage and elevated activity as measured with functional magnetic resonance imaging. J. Neurosci. 32, 12990–12998 (2012).

18. Emrich, S. M., Riggall, A. C., Larocque, J. J. & Postle, B. R. Distributed patterns of activity in sensory cortex reflect the precision of multiple items maintained in visual short-term memory. J. Neurosci. 33, 6516–6523 (2013).

19. Ester, E. F., Anderson, D. E., Serences, J. T. & Awh, E. A neural measure of precision in visual working memory. J. Cogn. Neurosci. 25, 754–761 (2013).

20. Lee, S.-H., Kravitz, D. J. & Baker, C. I. Goal-dependent dissociation of visual and prefrontal cortices during working memory. Nat. Neurosci. 16, 997–999 (2013).

21. Xing, Y., Ledgeway, T., McGraw, P. V. & Schluppeck, D. Decoding working memory of stimulus contrast in early visual cortex. J. Neurosci. 33, 10301–10311 (2013).

22. Albers, A. M., Kok, P., Toni, I., Dijkerman, H. C. & de Lange, F. P. Shared representations for working memory and mental imagery in early visual cortex. Curr. Biol. 23, 1427–1431 (2013).

23. Sprague, T. C., Ester, E. F. & Serences, J. T. Reconstructions of information in visual spatial working memory degrade with memory load. Curr. Biol. 24, 2174–2180 (2014).

24. Ester, E. F., Sprague, T. C. & Serences, J. T. Parietal and Frontal Cortex Encode Stimulus-Specific Mnemonic Representations during Visual Working Memory. Neuron 87, 893–905 (2015).

25. Sprague, T. C., Ester, E. F. & Serences, J. T. Restoring Latent Visual Working Memory Representations in Human Cortex. Neuron 91, 694–707 (2016).

26. Yu, Q. & Shim, W. M. Occipital, parietal, and frontal cortices selectively maintain task-relevant features of multi-feature objects in visual working memory. Neuroimage 157, 97–107 (2017).

27. Christophel, T. B., Klink, P. C., Spitzer, B., Roelfsema, P. R. & Haynes, J.-D. The Distributed Nature of Working Memory. Trends Cogn. Sci. 21, 111–124 (2017).

28. Rahmati, M., Saber, G. T. & Curtis, C. E. Population Dynamics of Early Visual Cortex during Working Memory. J. Cogn. Neurosci. 30, 219–233 (2018).

29. Christophel, T. B., Iamshchinina, P., Yan, C., Allefeld, C. & Haynes, J.-D. Cortical specialization for attended versus unattended working memory. Nat. Neurosci. 21, 494–496 (2018).

30. Lorenc, E. S., Sreenivasan, K. K., Nee, D. E., Vandenbroucke, A. R. E. & D’Esposito, M. Flexible Coding of Visual Working Memory Representations during Distraction. J. Neurosci. 38, 5267–5276 (2018).

31. Rademaker, R. L., Chunharas, C. & Serences, J. T. Coexisting representations of sensory and mnemonic information in human visual cortex. Nat. Neurosci. 22, 1336–1344 (2019).

32. Rahmati, M., DeSimone, K., Curtis, C. E. & Sreenivasan, K. K. Spatially Specific Working Memory Activity in the Human Superior Colliculus. J. Neurosci. 40, 9487–9495 (2020).

33. Brissenden, J. A., Tobyne, S. M., Halko, M. A. & Somers, D. C. Stimulus-Specific Visual Working Memory Representations in Human Cerebellar Lobule VIIb/VIIIa. J. Neurosci. 41, 1033–1045 (2021).

34. Tomko, G. J. & Crapper, D. R. Neuronal variability: non-stationary responses to identical visual stimuli. Brain Res. 79, 405–418 (1974).

35. Tolhurst, D. J., Movshon, J. A. & Dean, A. F. The statistical reliability of signals in single neurons in cat and monkey visual cortex. Vision Res. 23, 775–785 (1983).

36. Faisal, A. A., Selen, L. P. J. & Wolpert, D. M. Noise in the nervous system. Nat. Rev. Neurosci. 9, 292–303 (2008).

37. Foldiak, P. in Computation and Neural Systems (eds. Eeckman, F. H. & Bower, J.) (Springer Science & Business Media, 1993).

38. Sanger, T. D. Probability density estimation for the interpretation of neural population codes. J. Neurophysiol. 76, 2790–2793 (1996).

39. Zemel, R. S., Dayan, P. & Pouget, A. Probabilistic interpretation of population codes. Neural Comput. 10, 403–430 (1998).

40. Ma, W. J., Beck, J. M., Latham, P. E. & Pouget, A. Bayesian inference with probabilistic population codes. Nat. Neurosci. 9, 1432–1438 (2006).

41. Jazayeri, M. & Movshon, J. A. Optimal representation of sensory information by neural populations. Nat. Neurosci. 9, 690–696 (2006).

42. van Bergen, R. S., Ma, W. J., Pratte, M. S. & Jehee, J. F. M. Sensory uncertainty decoded from visual cortex predicts behavior. Nat. Neurosci. 18, 1728–1730 (2015).

43. van Bergen, R. S. & Jehee, J. F. M. Probabilistic Representation in Human Visual Cortex Reflects Uncertainty in Serial Decisions. J. Neurosci. 39, 8164–8176 (2019).

44. Walker, E. Y., Cotton, R. J., Ma, W. J. & Tolias, A. S. A neural basis of probabilistic computation in visual cortex. Nat. Neurosci. 23, 122–129 (2020).

45. van Bergen, R. S. & Jehee, J. F. M. TAFKAP: An improved method for probabilistic decoding of cortical activity. bioRxiv 2021.03.04.433946 (2021) doi:10.1101/2021.03.04.433946.

46. Hikosaka, O. & Wurtz, R. H. Visual and oculomotor functions of monkey substantia nigra pars reticulata. III. Memory-contingent visual and saccade responses. J. Neurophysiol. 49, 1268–1284 (1983).

47. Funahashi, S., Bruce, C. J. & Goldman-Rakic, P. S. Mnemonic coding of visual space in the monkey’s dorsolateral prefrontal cortex. J. Neurophysiol. 61, 331–349 (1989).

48. Mackey, W. E., Winawer, J. & Curtis, C. E. Visual field map clusters in human frontoparietal cortex. Elife 6, (2017).

49. Dumoulin, S. O. & Wandell, B. A. Population receptive field estimates in human visual cortex. Neuroimage 39, 647–660 (2008).

50. Hallenbeck, G. E., Sprague, T. C., Rahmati, M., Sreenivasan, K. K. & Curtis, C. E. Working Memory Representations in Visual Cortex Mediate the Effects of Distraction. Cold Spring Harbor Laboratory 2021.02.01.429259 (2021) doi:10.1101/2021.02.01.429259.

51. Kiani, R., Corthell, L. & Shadlen, M. N. Choice certainty is informed by both evidence and decision time. Neuron 84, 1329–1342 (2014).

52. Beck, J. M. et al.. Probabilistic population codes for Bayesian decision making. Neuron 60, 1142–1152 (2008).

53. Graf, A. B. A., Kohn, A., Jazayeri, M. & Movshon, J. A. Decoding the activity of neuronal populations in macaque primary visual cortex. Nat. Neurosci. 14, 239–245 (2011).

54. Berens, P. et al.. A fast and simple population code for orientation in primate V1. J. Neurosci. 32, 10618–10626 (2012).

55. Christophel, T. B., Hebart, M. N. & Haynes, J.-D. Decoding the contents of visual short-term memory from human visual and parietal cortex. J. Neurosci. 32, 12983–12989 (2012).

56. Brouwer, G. J. & Heeger, D. J. Decoding and reconstructing color from responses in human visual cortex. J. Neurosci. 29, 13992–14003 (2009).

57. Nienborg, H., Cohen, M. R. & Cumming, B. G. Decision-related activity in sensory neurons: correlations among neurons and with behavior. Annu. Rev. Neurosci. 35, 463–483 (2012).

58. Britten, K. H., Newsome, W. T., Shadlen, M. N., Celebrini, S. & Movshon, J. A. A relationship between behavioral choice and the visual responses of neurons in macaque MT. Vis. Neurosci. 13, 87–100 (1996).

59. Nienborg, H. & Cumming, B. G. Decision-related activity in sensory neurons reflects more than a neuron’s causal effect. Nature 459, 89–92 (2009).

60. Dodd, J. V., Krug, K., Cumming, B. G. & Parker, A. J. Perceptually bistable three-dimensional figures evoke high choice probabilities in cortical area MT. J. Neurosci. 21, 4809–4821 (2001).

61. Camarillo, L., Luna, R., Nácher, V. & Romo, R. Coding perceptual discrimination in the somatosensory thalamus. Proc. Natl. Acad. Sci. U. S. A. 109, 21093–21098 (2012).

62. Yu, X.-J., Dickman, J. D., DeAngelis, G. C. & Angelaki, D. E. Neuronal thresholds and choice-related activity of otolith afferent fibers during heading perception. Proc. Natl. Acad. Sci. U. S. A. 112, 6467–6472 (2015).

63. Goris, R. L. T., Ziemba, C. M., Stine, G. M., Simoncelli, E. P. & Movshon, J. A. Dissociation of Choice Formation and Choice-Correlated Activity in Macaque Visual Cortex. J. Neurosci. 37, 5195–5203 (2017).

64. Bang, D. & Fleming, S. M. Distinct encoding of decision confidence in human medial prefrontal cortex. Proc. Natl. Acad. Sci. U. S. A. 115, 6082–6087 (2018).

65. Lebreton, M., Abitbol, R., Daunizeau, J. & Pessiglione, M. Automatic integration of confidence in the brain valuation signal. Nat. Neurosci. 18, 1159–1167 (2015).

66. De Martino, B., Fleming, S. M., Garrett, N. & Dolan, R. J. Confidence in value-based choice. Nat. Neurosci. 16, 105–110 (2013).

67. Lee, S.-H. & Baker, C. I. Multi-Voxel Decoding and the Topography of Maintained Information During Visual Working Memory. Front. Syst. Neurosci. 10, 2 (2016).

68. Breedlove, J. L., St-Yves, G., Olman, C. A. & Naselaris, T. Generative Feedback Explains Distinct Brain Activity Codes for Seen and Mental Images. Curr. Biol. 30, 2211–2224.e6 (2020).

69. Favila, S. E., Kuhl, B. A. & Winawer, J. Perception and memory have distinct spatial tuning properties in human visual cortex. Cold Spring Harbor Laboratory 811331 (2020) doi:10.1101/811331.

70. Curtis, C. E. & D’Esposito, M. Persistent activity in the prefrontal cortex during working memory. Trends Cogn. Sci. 7, 415–423 (2003).

71. Sreenivasan, K. K., Curtis, C. E. & D’Esposito, M. Revisiting the role of persistent neural activity during working memory. Trends Cogn. Sci. 18, 82–89 (2014).

72. Blanke, O. et al.. Visual activity in the human frontal eye field. Neuroreport 10, 925–930 (1999).

73. Curtis, C. E. & Connolly, J. D. Saccade Preparation Signals in the Human Frontal and Parietal Cortices. Journal of Neurophysiology vol. 99 133–145 (2008).

74. Bruce, C. J. & Goldberg, M. E. Primate frontal eye fields. I. Single neurons discharging before saccades. J. Neurophysiol. 53, 603–635 (1985).

75. Sommer, M. A. & Wurtz, R. H. Frontal eye field sends delay activity related to movement, memory, and vision to the superior colliculus. J. Neurophysiol. 85, 1673–1685 (2001).

76. Armstrong, K. M., Chang, M. H. & Moore, T. Selection and maintenance of spatial information by frontal eye field neurons. J. Neurosci. 29, 15621–15629 (2009).

77. Moore, T. & Fallah, M. Control of eye movements and spatial attention. Proc. Natl. Acad. Sci. U. S. A. 98, 1273–1276 (2001).

78. Moore, T. & Armstrong, K. M. Selective gating of visual signals by microstimulation of frontal cortex. Nature 421, 370–373 (2003).

79. Bruce, C. J., Goldberg, M. E., Bushnell, M. C. & Stanton, G. B. Primate frontal eye fields. II. Physiological and anatomical correlates of electrically evoked eye movements. J. Neurophysiol. 54, 714–734 (1985).

80. Tehovnik, E. J., Sommer, M. A., Chou, I. H., Slocum, W. M. & Schiller, P. H. Eye fields in the frontal lobes of primates. Brain Res. Brain Res. Rev. 32, 413–448 (2000).

81. Wozny, D. R., Beierholm, U. R. & Shams, L. Probability matching as a computational strategy used in perception. PLoS Comput. Biol. 6, (2010).

82. Girshick, A. R., Landy, M. S. & Simoncelli, E. P. Cardinal rules: visual orientation perception reflects knowledge of environmental statistics. Nat. Neurosci. 14, 926–932 (2011).

83. Wei, X.-X. & Stocker, A. A. A Bayesian observer model constrained by efficient coding can explain ‘anti-Bayesian’ percepts. Nat. Neurosci. 18, 1509–1517 (2015).

84. Daneman, M. & Carpenter, P. A. Individual differences in working memory and reading. Journal of Verbal Learning and Verbal Behavior 19, 450–466 (1980).

85. Süß, H.-M., Oberauer, K., Wittmann, W. W., Wilhelm, O. & Schulze, R. Working-memory capacity explains reasoning ability—and a little bit more. Intelligence 30, 261–288 (2002).

86. Mackey, W. E. & Curtis, C. E. Distinct contributions by frontal and parietal cortices support working memory. Sci. Rep. 7, 6188 (2017).

87. Kay, K. N., Winawer, J., Mezer, A. & Wandell, B. A. Compressive spatial summation in human visual cortex. J. Neurophysiol. 110, 481–494 (2013).

88. Ledoit, O. & Wolf, M. A well-conditioned estimator for large-dimensional covariance matrices. J. Multivar. Anal. 88, 365–411 (2004).

89. Berens, P. CircStat: a MATLAB toolbox for circular statistics. J. Stat. Softw. 31, 1–21 (2009).

